# Molecular Pathophysiology of Cardiac Injury and Cardiac Microthrombi in Fatal COVID-19: Insights from Clinico-histopathologic and Single Nuclei RNA Sequencing Analyses

**DOI:** 10.1101/2021.07.27.453843

**Authors:** Nobuaki Fukuma, Michelle L. Hulke, Michael I. Brener, Stephanie Golob, Robert Zilinyi, Zhipeng Zhou, Christos Tzimas, Ilaria Russo, Claire McGroder, Ryan Pfeiffer, Alexander Chong, Geping Zhang, Daniel Burkhoff, Martin B. Leon, Mathew Maurer, Jeffrey W. Moses, Anne-Catrin Uhlemann, Hanina Hibshoosh, Nir Uriel, Matthias J. Szabolcs, Björn Redfors, Charles C. Marboe, Matthew R. Baldwin, Nathan R. Tucker, Emily J. Tsai

## Abstract

Cardiac injury is associated with critical COVID-19, yet its etiology remains debated. To elucidate the pathogenic mechanisms of COVID-19-associated cardiac injury, we conducted a single-center prospective cohort study of 69 COVID-19 decedents. Of six cardiac histopathologic features, microthrombi was the most commonly detected (n=48, 70%). We tested associations of cardiac microthrombi with biomarkers of inflammation, cardiac injury, and fibrinolysis and with in-hospital antiplatelet therapy, therapeutic anticoagulation, and corticosteroid treatment, while adjusting for multiple clinical factors, including COVID-19 therapies. Higher peak ESR and CRP during hospitalization were independently associated with higher odds of microthrombi. Using single nuclei RNA-sequence analysis, we discovered an enrichment of pro-thrombotic/anti-fibrinolytic, extracellular matrix remodeling, and immune-potentiating signaling amongst cardiac fibroblasts in microthrombi-positive COVID-19 hearts relative to microthrombi-negative COVID-19. Non-COVID-19 non-failing hearts were used as reference controls. Our cumulative findings identify the specific transcriptomic changes in cardiac fibroblasts as salient features of COVID-19-associated cardiac microthrombi.

---

In severe COVID-19, cardiac manifestations like arrhythmias,^1^ myocardial infarction,^2, 3^ acute heart failure,^4–6^ and cardiogenic shock^7^ are prevalent and associated with increased mortality.^8, 9^ Both direct and indirect mechanisms of COVID-19-associated cardiac injury have been postulated,^10^ yet details remain elusive. Early theories that COVID-19 cardiac injury reflected acute viral myocarditis have since been disproven by the rarity of cases that meet histologic criteria for myocarditis. Additionally, diverse cardiac histopathologic findings, including endothelial cell damage and inflammatory cell infiltrates without associated cardiac necrosis,^2, 11, 12^ have been described on autopsy of COVID-19 decedents but at variable prevalence. Subsequently, the detection of microvascular thrombi in the hearts,^13, 14^ lungs,^15–19^ liver,^20^ brain,^21^ and skin^22^ of COVID-19 patients have raised the possibility that, in severe cases, COVID-19 may be more akin to a systemic illness. Such reports of multi-organ microthrombi, along with a high incidence of venous thromboembolism in severe and critical COVID-19, have led investigators to hypothesize that severe SARS-CoV-2 infection promotes a hypercoagulable state, referred to as COVID-19-associated coagulopathy.^23–27^ However, neither empiric intermediate-dose prophylactic anticoagulation nor therapeutic anticoagulation for critical COVID-19 appear to reduce thrombosis, the need for organ support, or mortality.^28–30^ Emerging bench studies and clinical observations suggest that thrombosis in COVID-19 may be primarily immune-mediated and therefore targeting immunothrombosis may be essential to prevent or treat thrombotic complications.^31, 32^ Clinical, patient-oriented studies have yet to test this theory.

Whether cardiac microthrombi underlie acute cardiac injury in severe COVID-19 is unclear at this time. Moreover, the mechanistic details of how SARS-CoV-2 triggers immunothrombosis has not been determined. Prior COVID-19 cardiac autopsy series focused primarily on detecting acute viral myocarditis and SARS-CoV-2. Most were limited to small sample sizes (<50 persons)^11, 13, 14, 16, 17, 33–35^ and had long post-mortem intervals (PMI, *i.e.* the time between death and autopsy).^13, 33–36^ Shown to compromise tissue RNA integrity and yield,^37^ particularly that of the heart,^38^ long PMIs hindered the potential molecular analyses of cardiac tissue in prior COVID-19 autopsy studies. Moreover, the prevalence of cardiac microthrombi in fatal COVID-19 is not well-established.

To investigate the molecular pathophysiology of cardiac injury and microthrombi in fatal COVID-19, we instituted specific measures to reduce PMI and optimize autopsy specimen quality for immunohistology and single nuclei RNA sequencing (snRNA-seq). We examined both left and right ventricular cardiac tissue samples of COVID-19 decedents and determined the prevalence of acute cardiac histopathologic features attributable to SARS-CoV-2 infection, including cardiac microthrombi. We analyzed patient clinical data, ventricular viral load, and histopathologic features of each heart to identify clinico-histopathologic associations that could provide insight into the pathogenesis of COVID-19-associated cardiac injury and cardiac microthrombi. We conducted snRNA-seq analysis of a subset of COVID-19 cardiac tissue with and without microthrombi in order to determine the compositional and cell type-specific transcriptomic changes that may underlie the development of COVID-19-associated cardiac microthrombi.

## Results

### Patient Characteristics

Our study cohort of 69 laboratory-confirmed COVID-19 decedents had a mean age of 72±10 (range 38-97) years and 20 (29%) were women. Decedents were predominantly Hispanic (59%) or Black (16%). Hypertension, diabetes, and obesity were the commonest comorbidities. The median [IQR] time from hospitalization to death was 16 [4-36] days. The majority of patients had multiple organ failure with 59% supported on mechanical ventilation for 13 [2-34] days, 54% receiving vasoactive medication, and 22% receiving renal replacement therapy. Nearly 60% patients received corticosteroids; far fewer received tocilizumab (16%) or remdesivir (3%). Peak high-sensitivity troponin (hs-TnT) exceeded 99^th^ percentile reference limits in 93% of decedents; median peak hs-TnT was 99 [47-227] ng/L. D-dimer, ESR, and CRP were also markedly elevated (Table 1). All autopsies were performed with a median PMI of 20 [5.5-42.5] hours; 20 autopsies had PMI <8 hrs.

**Table 1.**
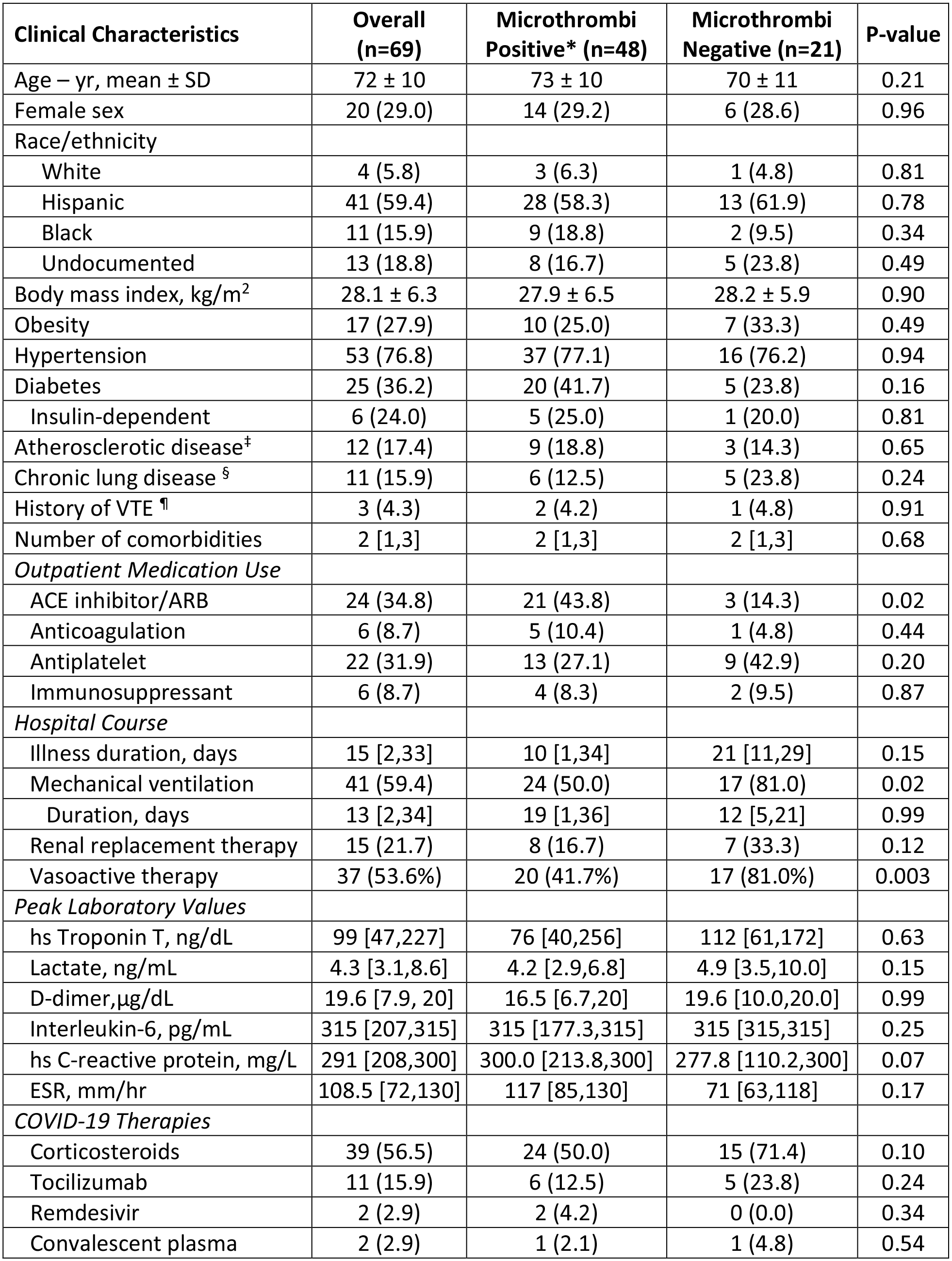
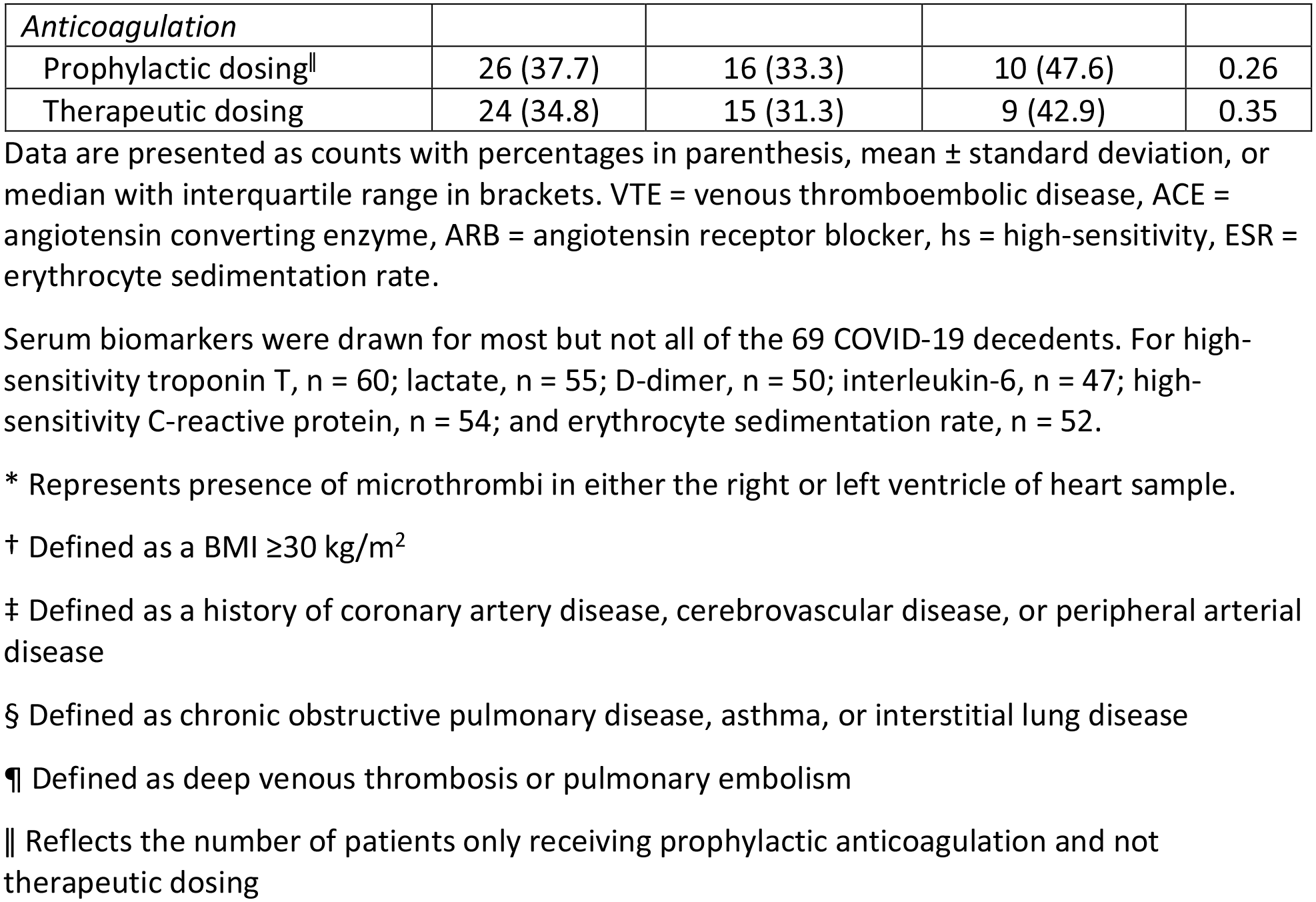
Study Cohort Characteristics.

### Cardiac Histopathology

Upon immunohistologic examination of both left and right ventricular tissue samples of the 69 hearts, we detected at least one of six acute cardiac histopathologic features in nearly all decedents (n=67, 97%): microthrombi (n=48, 70%), microvascular endothelial cell damage (n=25, 36%), scattered individual cardiomyocyte necrosis (n=25, 36%), focal cardiac necrosis without any adjacent inflammatory infiltrate (n=14, 20%), focal inflammatory infiltrates without associated cardiomyocyte injury (n=12, 17%), and focal myocarditis (n=4, 6%) (Fig. 1). Microthrombi were detected in serial sections by immunostaining for platelets (CD61). Microvascular endothelial cell damage was identified by complement protein C4d immunostaining, endothelial cell marker CD31 immunostaining, and by hematoxylin counterstain. Necrotic cardiomyocytes were likewise identified by C4d immunostaining and their characteristic branching, striated morphology appreciated on hematoxylin counterstain. Focal cardiac necrosis was defined as a confluence adjacent C4d+ cardiomyocytes occupying an area greater than 0.05mm^2^ but not exceeding 1cm^2^. Inflammatory infiltrates were detected by H&E staining. Most hearts (n=49, 71%) exhibited two or more of these acute histopathologic features, including microthrombi in 39 (80%) of such cases (Fig. 1G). Macroscopic and other histopathologic autopsy findings are summarized in Supplemental Table ST1.

**Figure 1.**
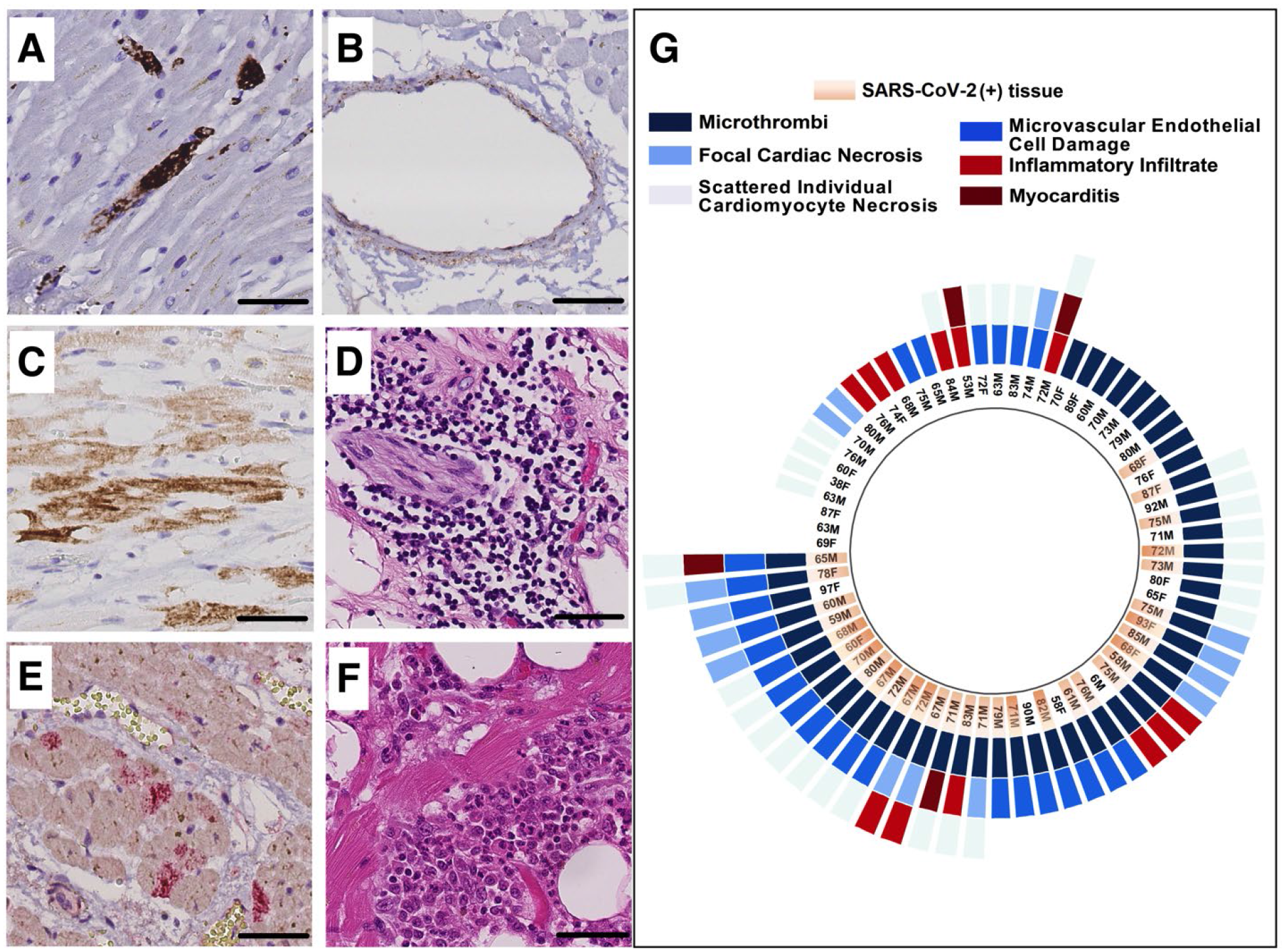
Acute cardiac histopathologic features in fatal COVID-19. **A.**Microthrombi, identified by CD61 immunostaining (DAB chromogen, brown). **B.** Small vessel endothelial cell damage, identified by C4d immunostaining (DAB chromogen, brown). **C.** Focal cardiac necrosis, identified by C4d+ cardiomyocytes (DAB chromogen, brown), in the absence of any adjacent inflammatory infiltrate. **D.** Inflammatory infiltrates on H&E. **E.** Scattered, isolated cardiomyocyte necrosis, identified by C4d+ staining (Alkaline Phosphatase chromogen, red). **F.** Focal myocarditis on H&E. Scale bars, 50µm. **G.** Radial plot of acute cardiac histopathologic features detected in the heart, present in either ventricle, of each COVID-19 decedent. Detection of SARS-CoV-2 in cardiac tissue, decedents’ age and sex are also shown.

In analyses adjusting for age and sex, microvascular endothelial damage was associated with cardiac microthrombi in the corresponding ventricle (OR 3.58, 95% CI 1.46-8.80). Cardiomyocyte necrosis, focal necrosis, inflammatory infiltrate, and myocarditis were not independently associated with cardiac microthrombi (Supplemental Table ST2).

### Ventricular Viral Load

We detected SARS-CoV-2 by RT-qPCR in 43 (62%) hearts. In analyses adjusting for age and sex, detectable SARS-CoV-2 in ventricular tissue was independently associated with microvascular endothelial cell damage in the corresponding ventricle (OR 2.36, 95% CI 1.04- 5.35). In contrast, ventricular viral load was not associated with the presence of microthrombi in the corresponding ventricle (Supplemental Table ST3).

### Biomarkers and clinical risk factors

In generalized additive logistic models (GAMs) adjusted for demographic, clinical, and COVID-19 treatment factors, peak hs-TnT and D-dimer were not associated with the predicted risk of cardiac microthrombi but that peak CRP was linearly associated with the predicted risk of cardiac microthrombi. ESR varied non-linearly with predicted risk of cardiac microthrombi, with plateaus of risk at levels <60 and >100 mm/hr (Fig. 2). In fully adjusted logistic regression models, every 20 mg/L change in CRP was associated with a 1.17 (95% CI 1.00-1.36) higher odds of cardiac microthrombi. Comparing patients in the third and fourth quartiles of ESR (107-126 and 130-169 mm/hr) to those in the first quartile (27-80 mm/hr), there was a 3.76 (95% CI 1.17- 12.04) and 6.65 (95% CI 1.53-28.79) higher odds of cardiac microthrombi, respectively (Supplemental Table ST4).

**Figure 2.**
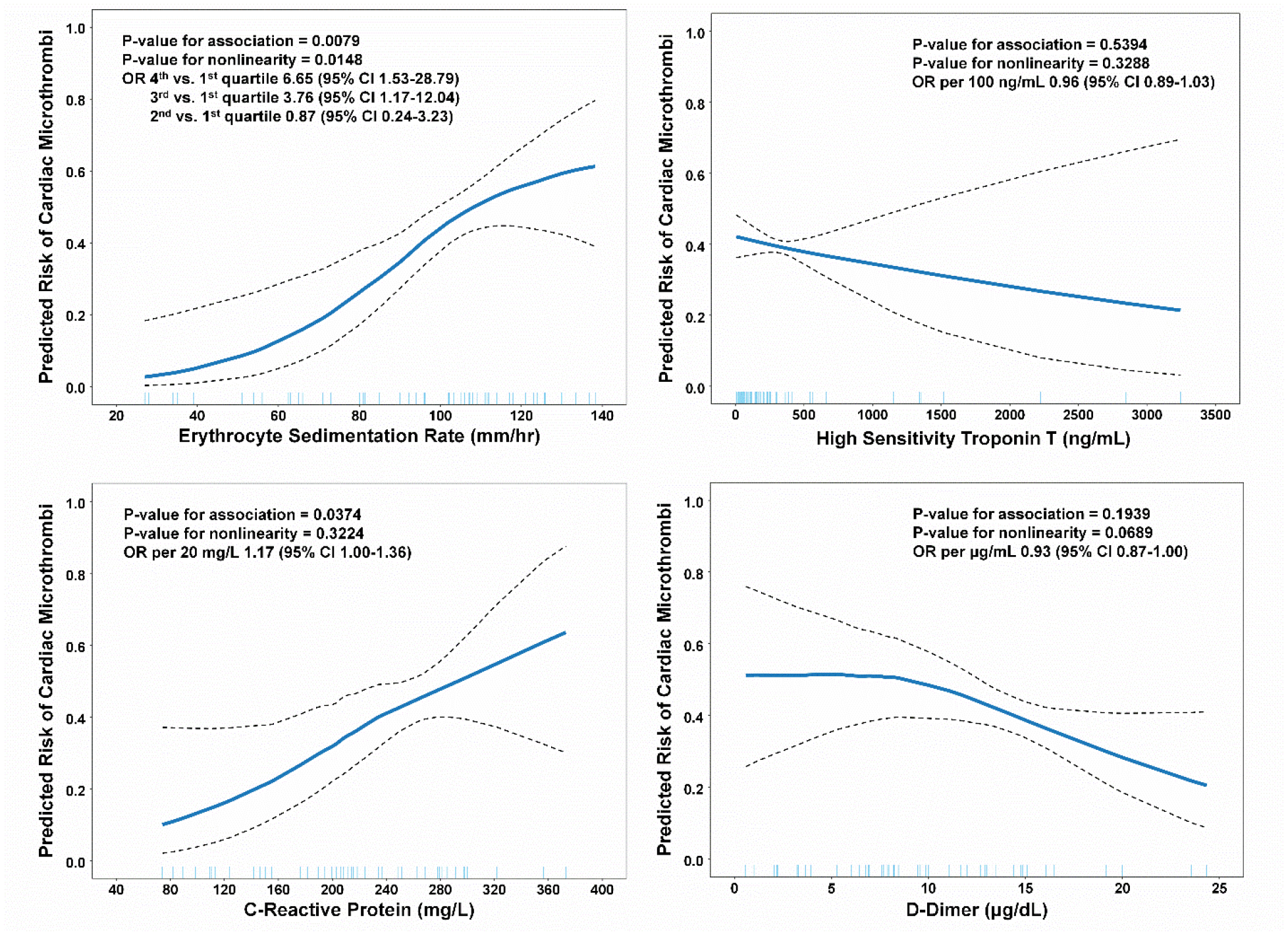
Adjusted generalized additive models (GAMs) of the association between serum biomarker peak values and cardiac microthrombi. We calculated a covariate balancing propensity score (CBPS) for each independent variable by regressing it on possible confounders; the resulting propensity score was used in the GAM model as a single covariable. The covariates used to calculate CBPS were: age, sex, race/ethnicity, body mass index (BMI), duration of COVID-19 illness, outpatient ACEi/ARB use, outpatient antiplatelet therapy, and inpatient administration of corticosteroids, remdesivir, interleukin-6 (IL-6) receptor antagonists, and therapeutic anticoagulation. OR, odds ratio.

In models adjusted for demographic, clinical, and treatment factors, use of outpatient antiplatelet therapy and in-hospital therapeutic anticoagulation were non-significantly associated with a lower odds of cardiac microthrombi, whereas in-hospital corticosteroid therapy was non-significantly associated with higher odds of cardiac microthrombi (Supplemental Fig. S1).

### Single Nuclei RNA Sequencing

To gain insight into the molecular pathophysiology of COVID-19-associated cardiac microthrombi, we performed snRNA-seq on mid-right ventricular free wall tissue from eight COVID-19 decedents with very low PMI (3.5 [2.8-5.0] hrs)--- four with and four without microthrombi (Supplemental Table ST5). No SARS-CoV-2 transcripts were detected in any sample--- an expected reflection of the known non-nuclear, cytosolic residence of SARS-CoV-2. We excluded one of the four microthrombi-positive samples from downstream analysis due to its low cellular complexity and high mitochondrial read count (Supplemental Table ST6). We compared data from the microthrombi-positive (13,068 cells) and microthrombi-negative (30,425 cells) COVID-19 samples with each other and with data from six non-failing hearts of the Human Cell Atlas (HCA)^39^ as the non-COVID-19 reference cohort (32,279 cells). Generated as part of a global effort to provide a non-failing reference map for analyses such as ours, the HCA heart reference has the potential to yield technically-driven effects. To minimize such potential, we reanalyzed the HCA reference from raw reads in parallel with our COVID-19 samples. Perfectly confounded, the HCA reference cohort could not be adjusted for any differences in sample collection, sample processing, or library generation. Thus, results from COVID-19 versus HCA reference comparison should be interpreted with this in mind. In contrast, microthrombi-positive and microthrombi-negative COVID-19 samples were collected and processed in parallel and therefore can be interpreted with greater confidence.

Principle components analysis of sample-level transcript counts clearly separated microthrombi-positive from microthrombi-negative COVID-19 samples and COVID-19 from non-COVID-19 reference samples (Supplemental Fig. S2). After stringent quality control measures (Supplemental Fig. S3), we analyzed the cellular diversity and transcriptional signatures of the aggregated snRNA-seq data, across all patient subsets (Fig. 3). We identified 12 distinct cell types (Fig. 3A), consistent with recent analyses of non-failing human myocardium.^39, 40^ Sensitive and specific transcriptional markers for each cell type are listed in Supplemental Table ST7.

**Figure 3.**
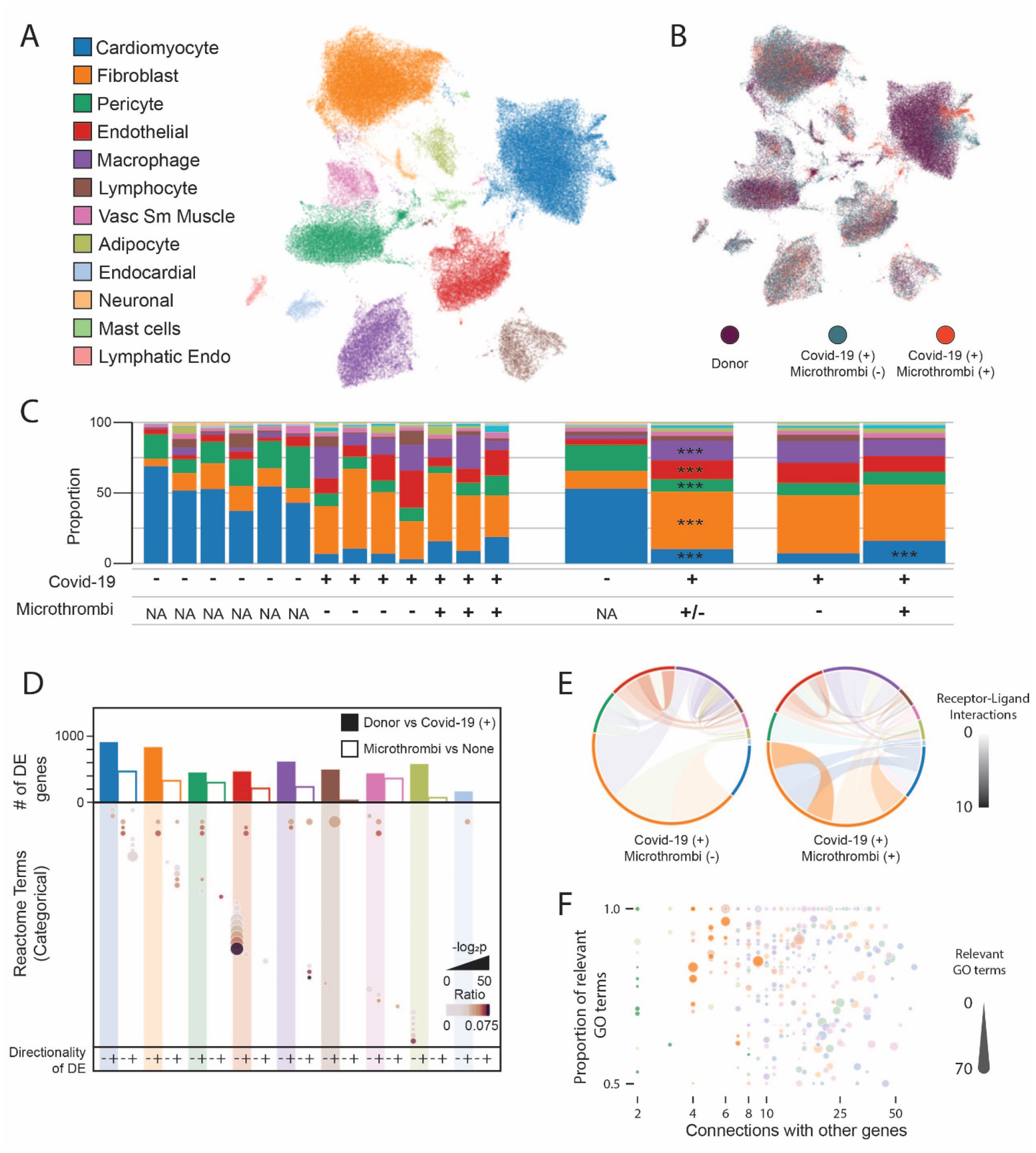
Single nucleus RNA sequencing of right ventricular tissue from COVID-19 autopsies. **A.**UMAP projection of 75,772 individual nuclei colored by cell type as determined from marker gene analysis. Colors observed within this legend are used throughout the figure. **B.** UMAP projection colored by sample type of origin. **C.** Compositional analysis for each cell type by sample (left panel) or condition (right panels). *** indicates a credible increase or decrease in cell proportion when compared to the referent dataset. **D.** Top bar plot displays the total number of DE genes at 0.25 log_2_ fold change (LFC) for COVID-19 versus non-COVID-19 control (solid bar) or microthrombi-positive versus microthrombi-negative in the COVID-19 (+) samples (open bar). Lower dot plot indicates the Reactome terms which are associated with DE genes in a given comparison and direction. Size and shade of the dot represent the adjusted P-value and ratio of Reactome genes present for a given term. **E.** CellPhoneDB analysis of cell-cell communication through analysis of receptor-ligand pair expression. Size of each outer section corresponds to the proportion of that cell type in the tissue. The opacity of each connection indicates the number of connections between cell types (paracrine and autocrine). Color of the connection indicates the cell type producing the ligand. **F.** Dot plot of Genewalk analysis identifying regulator genes for the ontology observed within the microthrombi versus none comparison within the COVID-19 (+) samples. Color corresponds to the cell type and size of the dot represents the number of significant GO terms affiliated with that gene. Dots with high opacity represent those from the 0.5 log_2_FC threshold for DE genes, while those with reduced opacity result from the more liberal 0.25 log_2_FC threshold.

To examine whether cell composition differs in COVID-19 versus non-COVID-19 reference hearts or in microthrombi-positive versus microthrombi-negative COVID-19 hearts, we quantified and compared relative proportions of each cell type in our snRNA-seq data, for each of the comparison groups (Fig. 3B,C). Any differences in cell composition could reflect either direct or indirect effects of SARS-CoV-2 infection, whereby the magnitude of difference may represent the relative role of or effect upon specific cell-types in COVID-19-associated cardiac injury and cardiac microthrombi. We identified credible differences in cell type proportions in COVID-19 versus reference samples--- decreases in cardiomyocytes (−2.33 log_2_ fold change (LFC)) and pericytes (−0.90 LFC) and increases in fibroblasts (1.67 LFC), endothelial cells (1.46 LFC), and macrophages (2.26 LFC) (Fig. 3C, Supplemental Table ST8). Microthrombi-negative COVID-19 samples exhibited greater loss of cardiomyocytes (0.60 LFC) than did the microthrombi-positive COVID-19 subset (Supplemental Table ST9).

We then identified the differentially expressed (DE) genes in the 9 most numerous cell types across subset comparisons of: a) COVID-19 versus non-COVID-19 reference control and b) microthrombi-positive versus microthrombi-negative COVID-19 (Supplemental Fig. S4 and Table ST10). We examined gene set enrichment in all DE genes for each subset comparison (Fig. 3D, Supplemental Table ST11). In COVID-19 versus reference control, multiple cell types shared several altered regulatory patterns. Upregulation of chromatin modifying enzymes suggest epigenetic changes in all but endothelial cells; and altered Rho-GTPases imply effects on cytoskeletal dynamics and cell motility in non-immune cells. We also uncovered cell type-specific features in COVID-19 samples compared to reference controls. In vascular endothelial cells, multiple pathways associated with IL-6 and interferon signaling were downregulated. By examining the DE genes that comprised these pathways, we found significant and marked downregulation of HLA class I proteins (HLA-A, B & C) and JAK/STAT family members---- a response that has been observed as a signature of viral immune evasion.^41^ In cardiomyocytes and vascular smooth muscle, pathways involved in striated and smooth muscle contraction were respectively downregulated, driven by genes encoding structural components of contractile apparati (*i.e.,* actinomyosin family members, troponins, and tropomyosins), signifying contractile dysfunction.

Cell type-specific signaling signatures were also noted in microthrombi-positive compared to microthrombi-negative COVID-19 specimens. In fibroblasts, extracellular matrix (ECM) pathways were markedly upregulated, including multiple genes that encode collagen and genes that encode proteins with dual anti-fibrinolytic/pro-thrombotic and innate immune response-related functions, such as *SERPINE1* and *THBS2*. In macrophages, cytokine signaling was upregulated. In cardiomyocytes, striated muscle contraction pathways were further downregulated, signifying more severe contractile dysfunction in microthrombi-positive COVID- 19 hearts.

To assess cell-cell communication, we then examined the expression of receptor-ligand pairs by each cell type population (Fig. 3E, Supplemental Table ST12). The number of cell-type specific signaling interactions involving fibroblasts was notably greater in microthrombi-positive compared to microthrombi-negative COVID-19 samples. With some autocrine and others paracrine in nature, the additional fibroblast signaling interactions in microthrombi-positive samples involved expression of multiple collagen types, the α11β1 integrin complex, and the platelet-derived growth factor D (PDGFD) and PDGF receptor complex.

We then used a network representation learning framework^42^ to identify regulator genes-- DE genes with the most connections to all other DE genes and to their gene ontologies. We searched for regulator genes that account for both the differences between COVID-19 and reference control (Supplemental Fig. S5, Supplemental Table ST13) and those between microthrombi-positive and microthrombi-negative COVID-19 samples (Fig. 3F). In sum, we identified 254 unique regulator genes, distributed across all cell types, that accounted for the differences between microthrombi-positive and microthrombi-negative COVID-19 samples (Supplemental Table ST13). Nearly a third of these regulator genes were shared by multiple cell types, indicating that a common cellular response is specific to COVID-19-associated cardiac microthrombi at the tissue level. To identify regulator genes with larger effect sizes, we examined genes with greater differential expression (≥0.5 LFC) between the microthrombi-positive and microthrombi-negative COVID-19 subsets. We uncovered 17 such large-effect regulator genes in fibroblasts and 9 in pericytes but none in any other cell types (Fig. 3F, Supplemental Tables ST13). Notably, *FGFR1* was one of the fibroblast-specific large-effect regulator genes, supporting the association of fibroblast proliferation and activation with the presence of microthrombi. Ontology terms underlying the connectivity of large-effect regulator genes revealed pro-thrombotic signaling associations including ECM deposition (collagen family members), promotion of platelet aggregation (*THBS1*), and inhibition of fibrinolysis (*STAT3*) in fibroblasts, further emphasizing that dysregulated fibroblast activity is a salient feature of COVID-19-associated cardiac microthrombi (Supplemental Fig. S6).

We intersected large-effect regulator genes with druggable genome annotations^43^ to prioritize targets for potential follow-up therapeutic investigation (Supplemental Table ST13). For COVID-19 versus reference control comparisons, 23 large-effect regulator genes were categorized in the druggable genome as Tier 1 (i.e., targeted by approved small molecules or late-stage drug candidates). Microthrombi-positive versus microthrombi-negative COVID-19 comparisons yielded four Tier 1 targets--- *PIP5K1A*, *STAT3*, and *VEGFA* in fibroblasts and *CSNK1A1* in pericytes.

Given the convergence of pro-fibrotic fibroblast signatures, we wished to specifically examine the activation state of fibroblasts in microthrombi-positive COVID-19 samples. We examined the expression of canonical markers of fibroblast activation, *FAP* and *POSTN,* in our snRNA-seq data. While we observed that the percentage of fibroblasts expressing these markers and the mean expression of these markers were higher in microthrombi-positive versus microthrombi-negative COVID-19 samples, they were not expressed broadly enough to be included in our wider DE model (Fig. 4A,B). To examine this further, we performed RT-qPCR on a subset of microthrombi-positive versus microthrombi-negative COVID-19 samples, including the samples used for snRNA-seq. Expression of *FAP* was significantly upregulated in the microthrombi-positive subset. While *POSTN* only trended toward higher expression levels in microthrombi-positive COVID-19 subset, this data still supports increased fibroblast activation at the tissue level (Fig. 4C).

**FIGURE 4.**
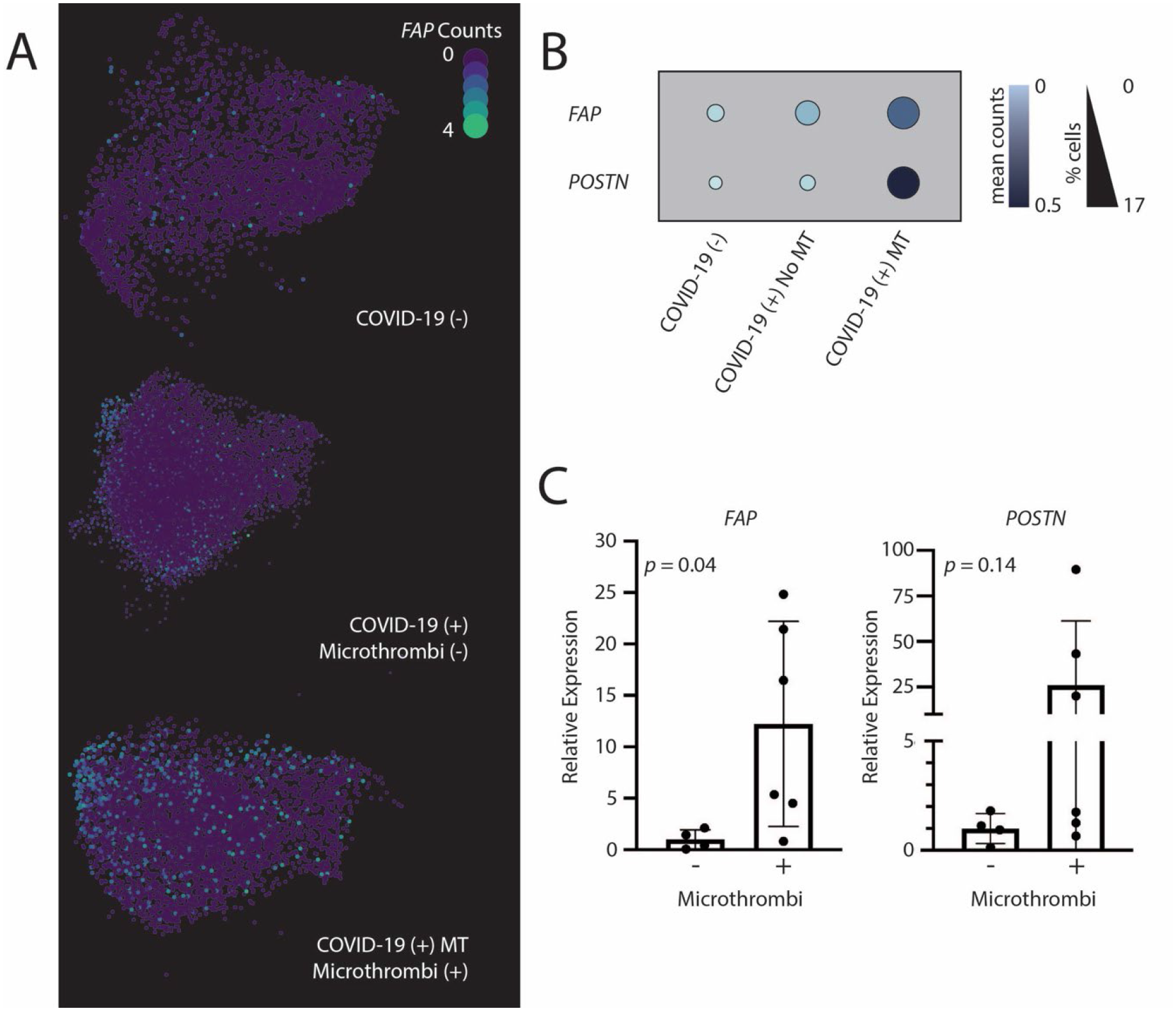
Analysis of fibroblasts activation in COVID-19 (+) samples with versus without microthrombi. A. UMAP projections of fibroblast cell populations. Colors correspond to the read counts for FAP in each cell. B. Dot plot displaying the expression of activated fibroblast markers *FAP* and *POSTN* in fibroblast clusters. Size and shade of the dot corresponds to the percentage of cells with non-zero counts and the mean counts across all cells, respectively. C. Relative cardiac tissue expression of *FAP* and *POSTN* (corrected to *RPS13*). P values shown for unpaired t test with Welch’s correction. See On-line Methods for details.

## Discussion

We conducted the first combined clinico-histopathologic and single cell analysis of a prospective COVID-19 cardiac autopsy cohort. We identified microthrombi as the predominant acute cardiac histopathologic feature in fatal COVID-19 and applied a multidisciplinary approach to investigating the pathophysiology of COVID-19-associated cardiac microthrombi. We found that biomarkers of systemic inflammation, but not those of fibrinolysis or cardiac injury, are independently associated with cardiac microthrombi, supporting an immunothrombotic mechanism of microthrombi formation. Our findings that anti-platelet and anticoagulation therapy have a non-significant association with a lower odds of cardiac microthrombi suggest that anticoagulation may be necessary but is insufficient in preventing and treating cardiac microthrombi in COVID-19. Our snRNA-seq analysis suggests that dysregulated cardiac fibroblasts expressing high levels of pro-thrombotic/anti-fibrinolytic, ECM-remodeling, and immune-potentiating genes are a salient feature of COVID-19-associated cardiac microthrombi. Thus, our cumulative findings provide new insights into the processes that may underlie COVID-19-associated immunothrombosis and cardiac microthrombi, with potential therapeutic implications for targeting cardiac fibroblast activity.

Our study is distinct from prior sc/snRNA-seq analyses of COVID-19 tissue specimens,^44, 45^ particularly that of COVID-19 cardiac samples, in three major aspects. First, by using immunohistology to systematically examine all ventricular samples, we could ascertain, without bias, that cardiac microthrombi is the predominant acute cardiac histopathologic feature in our fatal COVID-19 study population. This allowed us to optimize the translational significance of our snRNA-seq analyses. Moreover, the comparison between COVID-19 hearts with and without microthrombi provided an internal control for changes associated more generally with severe COVID-19. It enabled us to identify the cellular and transcriptomic changes that are most likely to be specific to cardiac microthrombi in severe COVID-19. Second, we examined associations of serum biomarkers with cardiac microthrombi, while controlling for multiple clinical covariates, including in-hospital therapies. Our findings suggest that an inflammatory process underlies cardiac microthrombi in COVID-19. Third, we also found an association, independent of age and sex, between cardiac microvascular endothelial cell damage and cardiac microthrombi in our cohort of COVID-19 decedents. This association is consistent with the understanding that immunothrombosis occurs in the context of endothelial cell dysfunction, immune cell activation, and platelet activation.^46^ That microthrombi in COVID- 19 is an immunothrombotic phenomenon has been widely assumed but, to date, unproven. The synergy of our clinical, histopathological, and molecular findings bolsters our conclusion that cardiac microthrombi in critical COVID-19 is indeed an immunothrombotic event, and not the result of disordered fibrinolysis and coagulation alone.

Additional strengths of our study include our cohort demographics and low PMI. Our study cohort consisted predominantly of Black and Hispanic decedents, reflecting their disproportionate share of COVID-19 deaths in the U.S. relative to their overall population.^47^ Yielding 20 COVID-19 autopsies with PMIs <8hrs out of our total cohort of 69, our low PMI initiative afforded us the ability to pursue snRNA-seq of cardiac tissue. Most prior COVID-19 autopsy studies of the heart reported PMIs of multiple days;^13, 33–36^ such delays compromise tissue RNA integrity of post-mortem tissue for molecular analysis.^37^

Consistent with the recent two studies of cardiac microthrombi in fatal COVID-19,^13, 14^ we found that the minority of COVID-19 autopsies had any focal cardiac necrosis. The infrequency of cardiac necrosis in fatal COVID-19^13, 14, 49^ contrasts with the high prevalence, magnitude, and mortality risk of cTn elevation among critically ill COVID-19 patients,^9, 48, 49^ even after controlling for renal dysfunction.^50, 51^ This discrepancy raises the question of what, if not cardiac necrosis, leads to cTn elevation in critical COVID-19. More specifically, what is the pathophysiologic sequelae of cardiac microthrombi, if not cardiomyocyte necrosis?

Our cell composition and single nuclei transcriptomic data demonstrate that SARS-CoV-2 indeed impacts cardiomyocytes, resulting in their cell death and dysfunction. The reduced proportion of cardiomyocytes in COVID-19 relative to reference control samples, along with the low prevalence of focal cardiac necrosis and the lack of association between cardiac microthrombi and cardiac necrosis in our overall COVID-19 cohort, suggests that the loss of cardiomyocytes is due to non-necrotic cell death. Additional non-necrotic causes of serum cTn elevation, such as increased cardiomyocyte membrane permeability and vesicular release from cardiomyocytes,^52, 53^ may also be involved but are beyond the scope of our current study. We found no association between ventricular SARS-CoV-2 viral load and peak serum hs-cTn-T, suggesting that acute cardiac injury is likely an indirect effect of COVID-19. Moreover, no evidence of SARS-CoV-2 infection of cardiomyocytes emerged from our snRNA-seq analysis, further pointing towards an indirect mechanism of cardiomyocyte injury. Our snRNA-seq data also puts forth the possibility that cardiomyocyte injury in COVID-19 may be attributable to upregulation of cytokine signaling by macrophages within the heart. Upregulated macrophage cytokine signaling may also lead to cardiac pericyte cell death in COVID-19, as a lower proportion of pericytes, without any transcriptomic evidence of SARS-CoV-2 infection, was also observed in COVID-19 samples versus reference controls. Further studies are needed to determine the details of such an indirect mechanism of cardiomyocyte and pericyte cell death in severe COVID-19.

Our foremost discovery, on snRNA-seq analysis, is the specific prothrombotic, anti-fibrinolytic, and immune activating signaling of cardiac fibroblasts in microthrombi-positive COVID-19 hearts. We identified pro-fibrotic responses in COVID-19 samples which can promote *in situ* microthrombi formation, particularly ECM deposition pathways that include secreted proteins with dual pro-thrombotic/anti-fibrinolytic and innate immune response functions. Moreover, the greater number of receptor-ligand interactions involving fibroblasts in microthrombi-positive versus microthrombi-negative COVID-19 samples highlights the significance of fibroblast signaling in the pathophysiology of COVID-19-associated cardiac microthrombi.

Whether these transcriptomic signatures of cardiac fibroblasts are the cause or consequence of cardiac microthrombi in critical COVID-19 requires additional mechanistic studies. Regardless, both scenarios have significant implications with regards to therapeutic intervention. If causal, fibroblast activation and signaling would be an important therapeutic target for mitigating microthrombi formation in acute COVID-19. If the transcriptomic signature of cardiac fibroblasts is a result of cardiac microthrombi, countering those specific pathways may be important to the prevention or treatment of chronic cardiac sequelae, such as cardiac fibrosis or microvascular disease, in patients who survive severe or critical COVID-19. In either case, our overall findings open the possibility that small molecule inhibition of proteins encoded by fibroblast regulator genes (*e.g*. *FGFR1, STAT3*) may be a novel approach to the clinical management of COVID-19 patients, be it in the acute or post-acute phase of severe illness. Such therapeutic promise warrants further mechanistic investigation into the role of fibroblasts in the pathophysiology of cardiac microthrombi and immunothrombosis in COVID-19.

### Limitations

Patients were hospitalized prior to FDA-approved therapies, and our initial local practice, as of April 15, 2020, was to reserve steroids for only the most critically-ill appearing patients. Therefore, the nearly statistically significant association of steroid use with increased odds of cardiac microthrombi may reflect some confounding by an unmeasured greater severity of illness. During the study period, testing for other respiratory viruses (i.e., influenza or adenovirus) was not routine. Reassuringly, a study of suspected acute myocarditis patients earlier in the pandemic offers evidence that viral co-infection occurs <5% of the time.^54^ Our observed associations between clinical variables and presence of cardiac microthrombi may be affected by unmeasured residual confounding and missingness, though we were able to adjust for several confounders via CBPS and we employed multiple imputation. The magnitude of the effect estimates may not be generalizable given our single-center study design.

While affording previously unattainable resolution of cell type-specific transcription, snRNA-seq analyses have technological limitations which are discussed extensively in the results and on-line methods as they arise. Importantly, SARS-CoV-2 resides in the cytosol, which is removed during sample processing for snRNA-seq. As such, we could not and did not observe SARS-CoV-2 transcripts in our analysis and must rely upon indirect evidence of viral infection of individual cell types in our analysis. Along similar lines, snRNA-seq analysis cannot capture cellular or transcriptomic data on anuclear cells, namely platelets. We utilized an externally developed non-COVID-19 reference control; expansive conclusions regarding COVID-19 versus non-COVID-19 transcriptional signatures should be interpreted conservatively. We provide complete analysis but limit our discussion to only the most distinct findings, focusing on the features observed in internally developed microthrombi-positive versus microthrombi-negative COVID-19 comparisons. Finally, our sample size for snRNA-seq is limited but comparable to most other snRNA-seq analyses of pediatric^55^ and adult^39, 56^ human hearts.

### Conclusions

We identified cardiac fibroblast activation as a salient feature of microthrombi-positive hearts in fatal COVID-19. Our observational study cannot determine whether fibroblast activity is causative of or in reaction to the formation of cardiac microthrombi. However, our snRNA-seq analysis revealed several expression signatures identifying a constellation of pro-thrombotic triggers at cell-type resolution. Regulator genes and pathways involved in platelet function and thrombosis in fibroblasts were prominent in microthrombi-positive versus microthrombi-negative COVID-19 samples, rendering it plausible that fibroblasts are integral to the propagation of cardiac microthrombi in fatal COVID-19. In summary, our findings imply that therapies targeted against fibroblast proliferation and activation, such as inhibitors of FGFR1 or STAT3, may be beneficial to severely ill COVID-19 patients and warrant further investigation.

## Acknowledgements

We are indebted to the families who provided next-of-kin consent for autopsy of their loved one. We also wish to thank the NYP-CUIMC clinicians who cared for COVID-19 patients and the Department of Pathology autopsy team members who worked around the clock during initial surge of the pandemic. Special thanks to Drs. Glen S. Markowitz, Donald W. Landry, and Ira A. Tabas for their leadership and support of the low PMI/rapid autopsy initiative; and to Bruce Spohler and Michael Gross for their contribution to the Columbia Cardiovascular COVID-19 Research Program (C3RP).

## Contributions

E.J.T. provided overall supervision. E.J.T. and N.R.T conceived and led the study. M.I.B., S.G., R.Z., and E.J.T. built the clinical database. C.C.M. and M.J.S. performed pathological review of tissues. C.C.M., M.J.S., I.R., N.F., and E.J.T. performed analysis of immunohistology. N.F., C.T., A.C., G.Z., N.R.T., M.H., and R.P. performed experiments. E.J.T., N.R.T., N.F., M.H., M.I.B., C.T., and I.R. performed analyses. M.R.B., B.R., Z.Z., and C.M. designed and performed statistical analyses of clinico-histological data. N.R.T., M.H., and R.P. prepared samples for single cell analysis. N.R.T. and M.H. designed and performed computational analysis of single cell data. E.J.T. and H.H. oversaw tissue collection. E.J.T., M.R.B., C.C.M., M.J.S., H.H., M.I.B., D.B., M.B.L., M.M., J.W.M., A.U., and N.U. provided clinical expertise. E.J.T., N.U. designed the I.R.B. protocols. N.F., M.H., M.I.B., M.R.B., N.R.T., and E.J.T wrote the manuscript. All authors reviewed and approved the final manuscript.

## Methods

### Study Design, Patient, and Clinical Data

We conducted a single-center prospective cohort study of hospitalized patients, age ≥18 years, diagnosed with SARS-CoV-2 infection via quantitative reverse transcription polymerase chain reaction (qRT-PCR) of nasopharyngeal swab as part of clinical care or, for persons under investigation who were suspected of COVID-19, by postmortem qRT-PCR. This cohort represents the first 69 consecutive qRT-PCR-confirmed COVID-19 autopsies performed at Columbia University Irving Medical Center (CUIMC, New York, NY) who died between March 26 and June 16, 2020. Next-of-kin provided informed consent for autopsy. Patient and autopsy data were manually abstracted from the electronic medical record, as approved by the CUIMC Institutional Review Board (IRB-AAS9835). Additional histologic and immunohistologic microscopy of ventricular tissue samples were performed for this research project. Per the Code of Federal Regulations 45 CFR 46.102, the research analysis of post-mortem tissue (IRB- AAT0135) was exempt from IRB evaluation.

For snRNA-seq analysis, we selected 8 COVID-19 cases, for which snap frozen ventricular tissue was collected at autopsy, with PMI<8hrs, based upon the presence of cardiac microthrombi and the absence of focal necrosis or myocarditis on microscopy in the ventricle (see Methods subsection *Microscopic Imaging* for details). We obtained the snRNA sequences of 6 non-COVID-19 organ donors from the Human Cell Atlas (HCA)^39^ and processed the data as our non-COVID-19 reference cohort. The HCA decedents were organ donors who died of brain death, with primary diagnoses of stroke, suicide, or trauma. They did not have hypertension, diabetes, nor any other pre-existing clinical cardiac disease. All organ donors had normal left ventricular ejection fraction (LVEF>50%).

### Low Post-Mortem Interval (PMI) Initiative

During the initial COVID-19 surge in New York City, the Columbia University Irving Medical Center (CUIMC) Departments of Pathology and Medicine implemented a multi-disciplinary initiative to lower PMI. Infographics, cloud-based resources, and multiple daily HIPAA-compliant text communications to frontline clinicians were used to support the low PMI initiative. Clinicians directly involved in the care of decedents obtained autopsy consent from next-of-kin at the time of death notification or shortly thereafter. Witnessed verbal consent was approved and accepted by NYP-CUIMC since all hospital visitations were prohibited during this period. Documentation of informed autopsy consent was protocolized for the electronic medical record; a copy of the document was sent to the decedents’ next-of-kin by electronic or postal mail. Timely transfer to the autopsy suite was coordinated immediately upon attainment of autopsy consent.

### Autopsies

Mid-free wall samples of the left (LV) and right ventricles (RV) were formalin-fixed, paraffin-embedded (FFPE), cut (4µm thick slices), and then stained with hematoxylin and eosin (H&E). Microscopic histology exams were performed by anatomic pathologists. We defined active myocarditis as myocyte degeneration or necrosis in the presence of adjacent inflammatory infiltrate.^57^ For autopsies with PMI <8hrs, additional ventricular samples were immediately snap frozen and stored at −80°C for snRNAseq analysis.

### Microscopic imaging and quantitative analysis of histopathologic findings

Four micron thick sections of left and right ventricular FFPE tissue blocks per decedent were cut and labeled using an automated staining platform (Leica Bond 3, Leica Biosystems, Buffalo Grove, IL) with the following monoclonal antibodies against: C4d (Leica Biosystems, BOND Ready-to-Use Primary Antibody, clone SP91) in order to detect cell damage or, in the case of cardiomyocytes, necrosis; CD61 (Cell Marque, prediluted primary antibody, clone EP65,) to identify platelet-rich microthrombi; and CD31 (Leica Biosystems, BOND Ready-to-Use Primary Antibody, clone 1A10) to verify endothelial cell identity. The target antigen was detected using either a DAB chromogen or Alkaline Phosphatase Red Chromogen. The latter was used to avoid interference from other types of brown pigment (e.g., hemosiderin or lipofuscin). All samples were counterstained with hematoxylin. As such, cardiomyocytes were identified by their characteristic rod-like morphology. Slides of left and right ventricular samples were also stained with hematoxylin and eosin. All slides were scanned using a Leica SCN400 slide scanner (Leica Scanner Console software v102.0.7.5). Analysis of all digital slides was performed independently, in parallel, by at least two blinded investigators, using ImageScope (Aperio/Leica Biosystems Imaging, Inc. CA USA).

Cardiomyocyte necrosis was evaluated for each ventricular sample. Focal cardiac necrosis was defined as a confluence of C4d-positive cardiomyocytes covering an area of at least 0.05 mm^2^ but not exceeding 1 cm^2^. Scattered, individual cardiomyocyte necrosis was defined as the presence of C4d-positive cardiomyocytes that were individually surrounded by only C4d-negative cardiomyocytes.

### Measurement of ventricular SARS-CoV-2 viral load

Total RNA was extracted from formalin-fixed paraffin-embedded (FFPE) tissue samples of left (LV) and right ventricles (RV) of all 69 heart, using the Quick-RNA FFPE Miniprep Kit (Zymo Research, Irvine, CA) according to the manufacturer’s instructions. RNA elution was performed with nuclease free double-distilled H_2_O at a final volume of 75 μl. We performed qRT-PCR, using primer/probe sets for the *N1* and *N2* regions of the SARS-CoV-2 nucleocapsid gene and for the human RNase P gene (*RP*) (Integrated DNA Technologies), as described previously.^58^ A standard curve of *N2* ranging from 10^1^-10^5^ viral copies was generated from the 2019-nCoV_N_Positive Control (Integrated DNA Technologies, Coralville, IA). Samples were considered positive for SARS-CoV-2 only if all three transcripts--- *N1*, *N2*, and *RP---* were detected.

For each quantitative reverse transcription-polymerase chain reaction (qRT-PCR), we used 5 μL of the total 75 μL of extracted RNA per ventricular tissue sample. Hence, the calculated viral load of each *N2* qRT-PCR reaction was multiplied by 15 to yield the ventricular viral load of the sample. To account for potential differences in ventricular tissue sample sizes, ventricular viral loads were normalized to *RP* expression using the Δ cycle threshold (Ct) method. Specifically, a normalization ratio for *RP* expression was calculated as 2^-ΔCt^, whereby ΔCt was the difference between each sample’s *RP* Ct and the mean *RP* Ct of all samples. Thus, the normalized viral load for each ventricular tissue sample was calculated by dividing the raw ventricular viral load by the normalization ratio.

### Measurement of canonical gene markers of fibroblast activation

Snap frozen RV samples of COVID-19 hearts (n=6 with microthrombi, n=4 without microthrombi) were each cut into five 5 micron thick pieces. These RV samples included all those analyzed by snRNA-seq as well as an additional 3 microthrombi-positive hearts. The samples were immersed in DNA/RNA Shield Reagent (Zymo) to preserve the genetic integrity of the samples at ambient temperatures and to inactivate any SARS-CoV-2 viral particles. We extracted total RNA from these samples using the RNeasy kit (QIAGEN) according to the manufacturer’s instructions. We performed one-step multiplex RT-qPCR using the Reliance One-step Multiplex Supermix (Bio-Rad), PrimePCR Probe Assays (FAP-FAM500, POSTN-HEX500, RPS13-Cy5500; Bio-Rad), and CFX96 Touch Real Time PCR Detection System (Bio-Rad).

### Single Nuclei RNA Sequencing Analysis

We isolated nuclei from snap frozen ventricular tissue, as previously described,^56^ and generated molecularly barcoded single nuclei emulsions (target 8000 nuclei per device) using the 10x Genomics Chromium Controller with v3.1 NextGEM Single Cell 3’ Kit. Libraries were constructed according to manufacturer’s protocols with few modifications (Supplemental Methods) and multiplexed for sequencing on HiSeq2500 or NovaSeqS1 flow cells.

### Data processing

All data processing was performed in the Terra Platform (app.terra.bio) unless otherwise noted. Postprocessing of data was accomplished using SCANPY v1.7.1.^59^ We obtained FASTQ files for non-Covid-19 donor hearts from the Human Cell Atlas’ Heart Atlas^39^ and processed them in parallel. Raw base call sequencing files were de-multiplexed; FASTQ files were generated using the 10x Genomics CellRanger 4.0.0 in the Cumulus workflow.^60^ To remove homopolymers (A30, T30, G30, and C30) and the template switch oligo sequence (CCCATGTACTCTGCGTTGATACCACTGCTT and its complement AAGCAGTGGTATCAACGCAGA GTACATGGG), reads were trimmed using Cutadapt (v2.8)^61^ with default parameters. Trimmed reads were aligned to the GRCh38 pre-mRNA human reference with SARS-CoV2 annotations (NC_045512). Count matrices were generated using CellRanger Count (v12).

We inspected each sample for mapping quality based on the number of mapped reads per cell, percentage of mapped mitochondrial reads (%mitochondrial reads), and the shape of the unique molecular identifier (UMI) decay curve. One sample (SID 69-RV) demonstrated high levels of mitochondrial reads and was removed from further analyses. The remaining samples were filtered using CellBender v2.0 ^62^ with default settings to remove the ambient RNA byproducts of nuclear isolation. From this, 108,016 cells remained.

Quality control was performed on the individual sample level. For each cell, we calculated the ratio of reads mapping to exonic regions to total mapped reads using Scrinvex (version 13) (https://github.com/getzlab/scrinvex). Cells with an exon ratio greater than the 75^th^ percentile + IQR range were removed from downstream analyses due to increased cytoplasmic transcripts (3,182 cells). We also excluded droplets which contained more than one nucleus as identified by Scrublet^63^ (10,866 cells). Samples were also filtered to remove cells with reads mapped to less than 200 genes (11,280 cells) and cells with greater than 5% mitochondrial reads (9,825 cells).

The gene list was filtered for highly variable genes (minimum mean 0.0125, maximum mean 3, minimum dispersion 0.5), using a subset of 6,255 genes for cell clustering. To account for variable complexity per nucleus, counts were normalized to 10,000 unique molecules per nucleus and logarithmized. Total read count and % mitochondrial reads were regressed out (Scanpy preprocessing *regress_out*); data was scaled to a maximum value of 10. Ischemic time for non-COVID-19 reference controls**Error! Hyperlink reference not valid.** was unavailable. Hence, we could not account for effects of ischemic time; results relating to hypoxic signaling should be interpreted with caution.

We calculated principal components from the highly variable gene subset (scanpy.tl.pca(adata, svd_solver=’arpack’)) and then corrected the normalized data for batch effects using Harmonypy (version 0.0.5)^64^ with each sample considered as a unique batch. We used the batch-corrected PCs to calculate neighbors (scanpy.pp.neighbors(adata, n_neighbors=10, n_pcs=40)) and generate a UMAP (scanpy.tl.umap(adata)). Cells were then clustered using leiden clustering (scanpy.tl.leiden(adata)) at a resolution of 0.225.

### Marker gene and cell type identification

Genes were ranked using a Wilcoxon rank sum test (scanpy *rank_genes_groups*) for each cell-type cluster versus all other cells; and log_2_-fold change (FC) and percentage of cells expressing each gene were calculated. Area under the receiver operator curve (AUC) scores were calculated for all genes within each cell cluster using SciKit Learn *roc_auc_score*. Genes were considered markers of a given cluster with an AUC score >0.7 or a log_2_-FC >0.6.

### Compositional analyses

The proportion of each cell type was compared across samples using scCODA (version 0.1.1).^65^ Briefly, we constructed a Markov-Chain Monte Carlo model with Hamiltonian Monte Carlo sampling using cell type proportions between conditions (non-COVID-19 reference control versus COVID-19, or COVID-19 microthrombi-positive versus COVID-19 microthrombi-negative). Credible interval differences in cell type proportions were determined using spike-and-slab inclusion probabilities. Importantly, the proportion of a given cell type in a sample is governed by our ability to liberate the nuclei equivalently from the tissue and our ability to successfully identify cells compared to empty droplets. The former may be affected by cell death or increased fibrosis which our sectioning protocol is designed to mitigate. The latter is a more challenging problem for single nucleus RNA sequencing, which is influenced by the relative transcriptional complexity of various cell types, making transcript rich cell types such as cardiomyocytes and fibroblasts easy to identify with transcript poor cells such as immune cells more apt to be assigned as an empty droplet. Use of a probabilistic cell calling mechanism in CellBender^62^ is used to overcome this challenge.

### Differential expression testing

Differentially expressed genes (DEGs) were calculated for each major cell cluster separately using the MAST^66^ pipeline. We constructed a Hurdle model (zlm(∼condition + ngenes + (1 | sample_ID), sca,method=’glmer’, ebayes = FALSE, strictConvergence = FALSE)) with the normalized reads based on the cellular detection rate, the set condition (either donor control versus COVID-19, or COVID-19 microthrombi-positive versus COVID-19 microthrombi-negative), and biological individual. Only genes with non-zero expression in at least 15% of all cells were included. DEGs were identified based on an Benjamini-Hochberg false discovery rate (FDR) adjusted P-value <0.05. Importantly, each droplet is contaminated by ambient RNA, which is imperfectly removed informatically by design. Therefore, DE results should be interpreted with caution, particularly when examining non-major cell types (e.g., intracardiac neurons, lymphatic endothelial, mast cells) and genes highly expressed in cells as numerous as cardiomyocytes.

Genes which serve as markers of another cluster (AUC ≥ 0.7) were blacklisted to exclude potential differential contamination by ambient RNA as a driver of such effect. Gene lists were also compared to the secreted protein list obtained from the Human Protein Atlas (proteinatlas.org). Full lists of DE genes, including those which were blacklisted, are contained in Supplemental Table ST3.

### Reactome pathway enrichment

For each major cell type, we performed gene pathway enrichment using Reactome^67^ (Pathway browser version 3.7, database release 75) on DEGs with log_2_-FC >0.25, separated into up or downregulated genes. We also calculated pathway enrichment using the Reactome pathway from GSEA Msigdb^68^ (MSigDB database v7.2). Pathways were considered enriched within a cell type if identified as such by both Reactome and GSEA Msigdb (BH FDR adjusted P- value of ≤ 0.05). Reactome terms for all comparisons are available in Supplemental Table ST11.

### Cell-Cell Communication

Cell-cell communication was tested with CellphoneDB (version2.1.7)^69^ on each sample separately using normalized count data for the 9 largest cell types. Afterward, significant interactions were aggregated between microthrombi-positive and microthrombi-negative samples. Briefly, CellphoneDB identifies and compares ligand-receptor interaction pairs between cell types and compares the observed interactions to the expected interactions of a null distribution generated from randomly permuted cell labels. Default parameters were used for the analysis (10% threshold for cells expressing ligands and receptors, p-value = 0.05, 1000 iterations for generation of null distribution, curated interactions list compiled by CellphoneDB from UniProt, Ensembl, PDB, IMEx consortium, and IUPHAR).

### Regulator Genes

To identify regulator genes that function within and likely drive gene networks within a biological context, we analyzed DEGs with a log_2_-FC ≥0.5 for each cell type using GeneWalk^42^ (direction of differential expression was not incorporated). GeneWalk builds biologically relevant networks from provided gene lists, connecting genes and GO terms, and compares the network to random networks. GeneWalk was used with default parameters and an FDR-corrected P-value of 0.1. The identified regulator genes were compared to a list of druggable genes.^43^ The full list of regulator genes identified is available in Supplemental Table ST13.

### Statistical Analysis

We tested associations between the presence of ventricular SARS-CoV-2 and histopathologic findings using logistic regression while adjusting for age and sex. We examined unadjusted associations of demographic, clinical, and laboratory characteristics with the presence of cardiac microthrombi in COVID-19 hearts using Student’s *t*, Mann-Whitney, or χ^2^ tests. For laboratory markers with greater than 5% missingness, we used multiple imputation in unadjusted tests of association. We created separate generalized additive logistic models (GAMs) with LOESS smoothers to test adjusted associations of the risk of cardiac microthrombi with plasma biomarkers of systemic inflammation, fibrinolysis, and cardiac injury. Due to the moderate cohort size and prevalence of microthrombi, we used covariant balanced propensity scores (CBPS) to adjust for potential confounders and precision variables, which included age, sex, race/ethnicity, body mass index (BMI), duration of COVID-19 illness (defined as the time interval in days between death and documented onset of symptoms consistent with COVID-19 or initial contact with health care system for symptoms consistent with COVID-19), outpatient ACEi/ARB use, outpatient antiplatelet therapy, and inpatient administration of corticosteroids, remdesivir, interleukin-6 (IL-6) receptor antagonists, and therapeutic anticoagulation. We estimated adjusted odds ratios using logistic regression models if there was no evidence of non-linearity in the GAMs. Since ESR had a non-linear association, we estimated odds ratios for each quartile increase in biomarker concentration. We also examined associations of treatments with cardiac microthrombi using logistic regression with CBPS to adjust for age, sex, race-ethnicity, BMI, and duration of COVID-19 illness, remdesivir, and IL-6 receptor antagonists. Statistical significance was defined by a two-sided p<0.05. Analyses were performed using SAS (v9.4, SAS Institute) and R(v3.5.1).

