## Supplementary material for "Molecular Pathophysiology of Cardiac Injury and Cardiac Microthrombi in Fatal COVID-19: Insights from Clinico-histopathologic and Single Nuclei RNA Sequencing Analyses": Data Supplement

From the Department of Medicine, Division of Cardiology (MIB, SG, RZ, CT, IR, NF, JWM, MM, MBL, DB, NU, and EJT), Division of Infectious Diseases (AC, AU), Division of Pulmonary, Allergy & Critical Care Medicine (CM, MRB), and Department of Pathology (GZ, HH, JEG, MJS, and CCM), Columbia University Irving Medical Center and the Cardiovascular Research Foundation (ZZ, BR, DB, JWM, and MBL) – all in New York City; St. Francis Hospital Heart Center (JWM) in Roslyn, NY; the Masonic Medical Research Institute (MLH, RP, and NRT) in Utica, NY; the Cardiovascular Disease Initiative, the Broad Institute of MIT and Harvard (NRT) in Cambridge, MA.

| Items | Pages |
|---|---|
| **Supplemental Tables** | |
| **Table ST1: Cardiac histopathology and ventricular viral load of COVID-19 decedents** | 3 |
| **Table ST2: Association between microthrombi and other acute histopathologic features at the ventricular level** | 4 |
| **Table ST3: Association between detectable SARS-CoV-2 and acute histopathologic features at the ventricular level** | 5 |
| **Table ST4: Association between cardiac microthrombi and ESR as a categorical variable** | 6 |
| **Table ST5: Clinical characteristics of COVID-19 (+) snRNAseq subsets** | 7 |
| **Table ST6: Sample level quality control metrics for snRNAseq** | 9 |
| **Table ST7: Marker genes by cell type for snRNAseq** | 10 |
| **Table ST8: Compositional analysis from snRNAseq using scCODA comparing COVID-19(+) vs COVID-19(-) reference samples** | 11 |
| **Table ST9: Compositional analysis from snRNAseq using scCODA comparing COVID-19(+) samples with vs without microthrombi** | 12 |
| **Table ST10: Differentially expressed genes identified by snRNAseq** | 13 |
| **Table ST11: Gene set enrichment analysis by cell type and condition using ReactomeDB** | 13 |
| **Table ST12: Cell-cell communication as assessed through CellphoneDB** | 13 |
| **Table ST13: Regulator genes identified by Genewalk for COVID-19 (+) vs (-) and microthrombi (+) vs microthrombi (-) COVID-19 subsets** | 13 |
| **Supplemental Figures** | |
| **Figure S1: Forest plot of the association between outpatient and in-hospital treatments and cardiac microthrombi** | 14 |
| **Figure S2: Principal components analysis of sample level transcript abundance** | 15 |
| **Figure S3: Identification and removal of low-quality cells in snRNAseq data** | 16 |
| **Figure S4: Volcano plots displaying differential expression results for presence of COVID-19 and for presence of microthrombi** | 17 |
| **Figure S5: Genewalk regulators for COVID-19 (+) versus COVID-19 (-) reference samples** | 18 |
| **Figure S6: Ontology analysis for select regulator genes identified by Genewalk** | 19 |

**Table ST1. Cardiac histopathology and ventricular viral load of COVID-19 decedents**

| COVID-19 | Overall (n=69) | Microthrombi-Positive* (n=48) | Microthrombi-Negative (n=21) |
|---|---|---|---|
| ***Ventricular Viral load*** | | | |
| Detectable viral load | 43 (62.3) | 31 (64.5) | 12 (57.1) |
| SARS-CoV-2 viral load* (copies) | 1055 [307, 13285] | 1594 [562,13285] | 387 [196, 21811] |
| ***Cardiac Autopsy Findings, n(%)*** | | | |
| Left ventricular hypertrophy | 45 (65.2) | 34 (70.8) | 11 (52.4) |
| Left ventricular dilation | 18 (26.1) | 16 (33.3)§ | 2 (9.5) |
| Right ventricular hypertrophy | 21 (30.4) | 16 (33.3) | 5 (23.8) |
| Right ventricular dilation | 37 (53.6) | 27 (56.3) | 10 (47.6) |
| Coronary atherosclerosis | 39 (56.5) | 26 (54.2) | 13 (61.9) |
| Myocardial infarction | 14 (20.3) | 10 (20.8) | 4 (19.0) |
| Thrombus† | 2 (2.9) | 2 (4.2) | 0 (0.0) |
| Interstitial edema | 5 (7.2) | 3 (6.3) | 2 (9.5) |
| Perivascular fibrosis | 23 (33.3) | 16 (33.3) | 7 (33.3) |
| Interstitial fibrosis | 27 (39.1) | 21 (43.8) | 6 (28.6) |
| Wavy myocytes | 7 (10.1) | 3 (6.3) | 4 (19.0) |
| Contraction bands | 5 (7.2) | 3 (6.3) | 2 (9.5) |
| Pericardial findings | 12 (17.4) | 8 (16.7) | 4 (19.0) |
| ***Cardiac Histopathologic Findings‡, n(%)*** | | | |
| Microvascular endothelial cell damage | 25 (36.2) | 19 (39.6)§ | 6 (28.6) |
| Scattered individual cardiomyocyte necrosis | 25 (36.2) | 16 (33.3) | 9 (42.8) |
| Focal cardiac necrosis | 14 (20.3) | 11 (22.9) | 3 (14.3) |
| Focal inflammatory infiltrate | 12 (17.4) | 6 (12.5) | 6 (28.6) |
| Focal myocarditis | 4 (5.8) | 2 (4.2) | 2 (9.5) |
| ***Pulmonary Autopsy Findings, n(%)*** | | | |
| Diffuse alveolar damage | 38 (55.1) | 27 (56.3) | 11 (52.4) |
| Pulmonary artery thrombosis | 10 (14.5) | 7 (14.6) | 3 (14.3) |
| Pulmonary microvascular thrombi | 42 (60.9) | 31 (64.6) | 11 (52.4) |

Data are presented as counts with percentages in parenthesis and median with interquartile range in brackets.

* Based upon the higher value detected in either the left or right ventricle of decedent

† Intraventricular, intra-atrial, or epicardial coronary arterial thrombus

‡ Based upon detection in either left or right ventricle of decedent

§ $P<0.05$

**Table ST2. Association between microthrombi and other acute histopathologic features at the ventricular level.**

| Dependent Variable | Independent Variable | Univariate Model | | Multivariate Model | |
|---|---|---|---|---|---|
| | | OR (95% CI) | p | OR (95% CI) | p |
| Microthrombi | Microvascular endothelial cell damage | 3.44 (1.42, 8.32) | 0.006 | 3.58 (1.46, 8.80) | 0.005 |
| Scattered individual necrotic cardiomyocytes | Microthrombi | 1.00 (0.46, 2.20) | 1.00 | 1.05 (0.47, 2.34) | 0.90 |
| Focal cardiac necrosis | Microthrombi | 1.23(0.37, 4.14) | 0.74 | 1.03 (0.30, 3.56) | 0.96 |
| Focal inflammatory infiltrate | Microthrombi | 0.43 (0.14, 1.35) | 0.15 | 0.39 (0.12, 1.25) | 0.11 |
| Focal myocarditis | Microthrombi | 1.27 (0.21, 7.85) | 0.80 | 1.30 (0.20, 8.52) | 0.78 |

All histopathologic features listed above were identified on immunohistologic microscopy with the exception of focal inflammatory infiltrate and myocarditis, which were identified by H&E staining. See On-line Methods for details.Multivariate model adjusted with age and sex.

**Table ST3. Association between detectable SARS-CoV-2 and acute histopathologic features at the ventricular level**

| Histopathologic Feature | Univariate Model | | Multivariate Model | |
|---|---|---|---|---|
| | OR (95% CI) | p | OR (95% CI) | p |
| Microthrombi | 1.52 (0.78, 2.99) | 0.22 | 1.62 (0.81, 3.22) | 0.17 |
| Microvascular endothelial cell damage | 2.24 (1.00, 5.03) | 0.05 | 2.36 (1.04, 5.35) | 0.04 |
| Scattered individual necrotic cardiomyocytes | 1.42 (0.66, 3.04) | 0.37 | 1.43 (0.67, 3.04) | 0.35 |
| Focal cardiac necrosis | 0.40 (0.12, 1.27) | 0.12 | 0.42 (0.13, 1.36) | 0.15 |
| Focal inflammatory infiltrate or myocarditis | 0.30 (0.08, 1.03) | 0.06 | 0.28 (0.08, 1.04) | 0.06 |

All histopathologic features listed above were identified on immunohistologic microscopy with the exception of focal inflammatory infiltrate or myocarditis, which was identified by H&E staining. See On-line Methods for details. Multivariate model is adjusted with age and sex.

**Table ST4. Association between cardiac microthrombi and ESR as a categorical variable.**

| ESR | | Unadjusted OR (95% CI) | P-value | Adjusted OR (95% CI) | P-Value |
|---|---|---|---|---|---|
| **Quartile** | **mm/hr** | | | | |
| 1st | 27.0-80.0 | *Reference* | | | |
| 2nd | 81.0-107.1 | 1.70 (0.65-4.50) | 0.280 | 0.87 (0.24-3.23) | 0.840 |
| 3rd | 107.2-126.0 | 2.78 (1.08-7.17) | 0.034 | 3.76 (1.17-12.04) | 0.026 |
| 4th | 130.0-169.1 | 1.95 (0.77-4.91) | 0.160 | 6.65 (1.53-28.79) | 0.011 |

Logistic regression model was adjusted for possible confounders by calculating a covariate balancing propensity score (CBPS) and using it as a single covariable. The covariates used to calculate CBPS were: age, sex, race/ethnicity, body mass index, duration of Covid-19 illness, outpatient ACEi/ARB use, outpatient antiplatelet therapy, and inpatient administration of corticosteroids, remdesivir, interleukin-6 (IL-6) receptor antagonists, and therapeutic anticoagulation.

ESR = Erythrocyte sedimentation rate, OR = odds ratio, CI = confidence interval

**Table ST5. Clinical characteristics of COVID-19 (+) snRNAseq subset**

| | Microthrombi (+) * | | | Microthrombi (-) | | | |
|---|---|---|---|---|---|---|---|
| *Study ID* | *05* | *39* | *61* | *19* | *45* | *51* | *66* |
| ***Baseline Characteristics*** | | | | | | | |
| **Age – yr** | 83 | 71 | 58 | 68 | 65 | 63 | 69 |
| **Sex** | Male | Male | Male | Male | Male | Male | Female |
| **Race/ethnicity** | Hispanic | Hispanic | Hispanic | Hispanic | Hispanic | Hispanic | n/a |
| **Body mass index – kg/m²** | 24.0 | 34.7 | 28.5 | 32.0 | 29.0 | 34.5 | 23.0 |
| **Obesity†** | · | ✓ | · | ✓ | · | ✓ | · |
| **Hypertension** | ✓ | ✓ | · | ✓ | ✓ | ✓ | ✓ |
| **Diabetes** | · | ✓ | ✓ | · | · | · | · |
| **Insulin-dependent** | · | · | ✓ | · | · | · | · |
| **Atherosclerotic disease‡** | · | · | · | · | · | ✓ | · |
| **Chronic lung disease§** | · | · | · | · | ✓ | · | ✓ |
| **History of VTE¶** | · | · | · | · | · | · | ✓ |
| **Number of Comorbidities** | 1 | 2 | 1 | 1 | 2 | 2 | 3 |
| ***Outpatient Medication Use*** | | | | | | | |
| **ACE inhibitor/ARB** | · | ✓ | · | · | · | · | · |
| **Anticoagulation** | · | · | · | · | · | · | · |
| **Antiplatelet** | ✓ | · | · | · | ✓ | · | · |
| **Immunosuppressant** | · | · | · | · | ✓ | · | · |
| ***Clinical Course*** | | | | | | | |
| **Duration of illness – days** | 9 | 26 | 57 | 26 | 24 | 21 | 40 |
| **Mechanical ventilation** | · | ✓ | ✓ | ✓ | ✓ | ✓ | · |
| **Duration – days** | · | 0 | 57 | 21 | 7 | 9 | · |
| **Renal replacement therapy** | · | · | ✓ | · | ✓ | · | · |
| **Vasoactive support** | · | · | ✓ | ✓ | ✓ | ✓ | · |
| ***Laboratory Studies, peak values*** | | | | | | | |
| **hs Troponin T, ng/dL** | 410 | 24 | 292 | 542 | 212 | 30 | 61 |
| **Lactate, ng/mL** | 1.7 | 4.2 | 4.5 | 3.1 | 10.1 | 11.1 | 3.9 |
| **D-dimer, μg/dL** | 1.01 | 5.30 | 20.00 | 20.00 | 20.00 | 20.00 | 10.00 |
| **Interleukin-6, pg/mL** | 207.0 | 315.0 | 315.0 | 315.0 | 315.0 | 273.0 | 108.0 |
| **hs C-reactive protein, mg/L** | 124 | 300 | 300 | 278 | 234 | 109 | 278 |
| **ESR, mm/hr** | 35 | 109 | 130 | 63 | 39 | 27 | 130 |
| ***Covid-19 Therapies*** | | | | | | | |
| **Corticosteroids** | · | ✓ | ✓ | ✓ | ✓ | ✓ | ✓ |
| **Tocilizumab** | ✓ | ✓ | · | ✓ | ✓ | · | · |
| **Remdesivir** | · | · | · | · | · | · | · |
| **Convalescent plasma** | · | · | · | · | ✓ | · | · |
| ***Anticoagulation*** | | | | | | | |
| **Prophylactic dosing** | ✓ | ✓ | ✓ | · | ✓ | ✓ | · |
| **Therapeutic dosing** | · | · | · | ✓ | · | · | ✓ |

n/a=not available (undocumented)

✓ Represents presence of trait/therapy/finding

· Represents absence of trait/therapy/finding

VTE = venous thromboembolic disease

ACE = angiotensin converting enzyme

ARB = angiotensin receptor blocker

hs = high-sensitivity

ESR = erythrocyte sedimentation rate

* Represents microthrombi detected on immunohistochemical analysis of right ventricle

† Defined as a body mass index (BMI) ≥30 $kg/m^2$

‡ Defined as a history of coronary artery disease, cerebrovascular disease, or peripheral arterial disease

§ Defined as chronic obstructive pulmonary disease, asthma, or interstitial lung disease

¶ Defined as deep venous thrombosis or pulmonary embolism

‖ Reflects the number of patients only receiving prophylactic anticoagulation and not therapeutic dosing

**Table ST6. Sample level quality control metrics for snRNAseq**

| Study ID | Estimated Number of Cells | Mean Reads per Cell | Median Genes per Cell | Number of Reads | Total Genes Detected | Median UMI Counts per Cell | Microthrombi | Included in Analysis | Post QC Number of Cells |
|---|---|---|---|---|---|---|---|---|---|
| **05** | 9,575 | 10,223 | 791 | 97,892,749 | 28,791 | 1,122 | Positive | TRUE | 7208 |
| **39** | 3,497 | 56,326 | 758 | 196,972,045 | 26,402 | 1,285 | Positive | TRUE | 1846 |
| **51** | 9,886 | 21,626 | 1,910 | 213,804,137 | 35,115 | 3,979 | Negative | TRUE | 5787 |
| **19** | 14,427 | 34,978 | 2,072 | 504,630,120 | 38,265 | 3,735 | Negative | TRUE | 9947 |
| **45** | 9,683 | 31,419 | 1,670 | 304,235,656 | 35,936 | 3,092 | Negative | TRUE | 6851 |
| **66** | 9,150 | 11,807 | 1,607 | 108,041,509 | 33,527 | 3,059 | Negative | TRUE | 7840 |
| **61** | 5,412 | 15,253 | 1,158 | 82,552,819 | 31,457 | 1,871 | Positive | TRUE | 4014 |
| **69** | 3,905 | 13,434 | 266 | 52,460,970 | 26,291 | 416 | Positive | FALSE | 0 |

Study ID: COVID-19(+) sample ID
Estimated Number of Cells: Number of droplets called as cells from CellRanger pipeline
Mean Reads per Cell: Average number of mapped reads per cell
Median Genes per Cell: Median number of genes detected in cells
Number of Reads: Total number of reads with multiplexing index matching a given sample
Total Genes Detected: Total unique gene IDs detected across all cells
Median UMI Counts per Cell: Median number of unique transcript molecules detected per cell
Microthrombi: Detection of cardiac microthrombi by immunohistochemistry (CD61 staining of corresponding ventricular tissue)
Included in Analysis: Use of this sample in downstream analysis pipeline
Post-QC Number of Cells: Number of cells retained following filtering for aberrant mitochondrial reads, intron/exon ratio, and doublet score

**Table ST7 is attached as an excel spreadsheet**.

**Table ST8. Compositional analysis from snRNAseq using scCODA comparing COVID-19 (+) vs COVID-19 (-) reference samples**

| **Cell Type** | **$log_2$-FC Ref1** | **Inclusion Probability Ref1** | **Credible Ref1** | **$log_2$-FC Ref2** | **Inclusion Probability Ref2** | **Credible Ref2** |
|---|---|---|---|---|---|---|
| Cardiomyocyte | 2.33904 | 1 | TRUE | 2.344553 | 1 | TRUE |
| Fibroblast | -1.67547 | 0.962867 | TRUE | -1.656135 | 0.963267 | TRUE |
| Pericyte | 0.898065 | 0.962333 | TRUE | 0.902993 | 0.9368 | TRUE |
| Endothelial | -1.461012 | 0.831733 | TRUE | -1.465264 | 0.855 | TRUE |
| Macrophage | -2.257872 | 0.998267 | TRUE | -2.26632 | 1 | TRUE |
| Lymphocyte | -0.381312 | 0.3492 | FALSE | -0.38268 | 0 | FALSE |
| Smooth Muscle | -0.381312 | 0 | FALSE | -0.38268 | 0.4374 | FALSE |
| Adipocyte | -0.381312 | 0.389667 | FALSE | -0.38268 | 0.5376 | FALSE |
| Endocardial | -0.381312 | 0.438 | FALSE | -0.38268 | 0.546 | FALSE |
| Neuronal | -0.381312 | 0.378867 | FALSE | -0.38268 | 0.3857 | FALSE |
| MAST | -0.381312 | 0.586333 | FALSE | -0.38268 | 0.560333 | FALSE |
| Lymphatic Endothelial | -0.381312 | 0.562867 | FALSE | -0.38268 | 0.602733 | FALSE |

**Ref1 =** Smooth Muscle
**Ref2 =** Lymphocyte

**Cell Type** Identified cell type from marker genes
**$log_2$-FC** $Log_2$ fold change of cell type in COVID-19(+) samples compared to cell type in COVID-19(-) reference controls
**Inclusion Probability** Spike-and-slab inclusion probability
**Credible** True if inclusion probability is above the Spike-and-slab threshold

**Table ST9. Compositional analysis from snRNAseq using scCODA comparing COVID-19 (+) samples with vs without microthrombi**

| Cell Type | $log_2$-FC Ref1 | Inclusion Probability Ref1 | Credible Ref1 | $log_2$-FC Ref2 | Inclusion Probability Ref2 | Credible Ref2 |
|---|---|---|---|---|---|---|
| Cardiomyocyte | -0.600618 | 0.664333 | TRUE | -0.65457 | 0.685267 | TRUE |
| Fibroblast | 0.061448 | 0.392867 | FALSE | 0.06737 | 0.383867 | FALSE |
| Pericyte | 0.061448 | 0.472667 | FALSE | 0.06737 | 0.418067 | FALSE |
| Endothelial | 0.061448 | 0.440867 | FALSE | 0.06737 | 0.438333 | FALSE |
| Macrophage | 0.061448 | 0.452933 | FALSE | 0.06737 | 0 | FALSE |
| Lymphocyte | 0.061448 | 0.4988 | FALSE | 0.06737 | 0.524533 | FALSE |
| Smooth Muscle | 0.061448 | 0 | FALSE | 0.06737 | 0.476 | FALSE |
| Adipocyte | 0.061448 | 0.446267 | FALSE | 0.06737 | 0.494933 | FALSE |
| Endocardial | 0.061448 | 0.508867 | FALSE | 0.06737 | 0.521533 | FALSE |
| Neuronal | 0.061448 | 0.469 | FALSE | 0.06737 | 0.5274 | FALSE |
| MAST | 0.061448 | 0.533133 | FALSE | 0.06737 | 0.4994 | FALSE |
| Lymphatic Endothelial | 0.061448 | 0.524267 | FALSE | 0.06737 | 0.479067 | FALSE |

**Ref1 = Smooth Muscle**
**Ref2 = Macrophage**

| | |
|---|---|
| **Cell Type** | Identified cell type from marker genes |
| **$log_2$-FC** | $Log_2$ fold change in cell types in microthrombi(+) compared to microthrombi(-) COVID-19(+) samples |
| **Inclusion Probability** | Spike-and-slab inclusion probability |
| **Credible** | True if inclusion probability is above the Spike-and-slab threshold |

**Tables ST10-ST13 are attached as excel spreadsheets**

**Figure S1. Forest plot of the association between outpatient and in-hospital treatments and cardiac microthrombi.** Logistic regression models were adjusted for possible confounders by calculating a covariate balanced propensity score (CBPS) for each model and using it as a single covariable. The covariates used to calculate CBPS were: age, sex, race/ethnicity, body mass index (BMI), duration of COVID-19 illness, outpatient ACEi/ARB use, outpatient antiplatelet therapy, and inpatient administration of corticosteroids, remdesivir, interleukin-6 (IL-6) receptor antagonists, and therapeutic anticoagulation. For variables that were also listed as a covariate, the redundant covariate was excluded from the respective CBPS for that model.

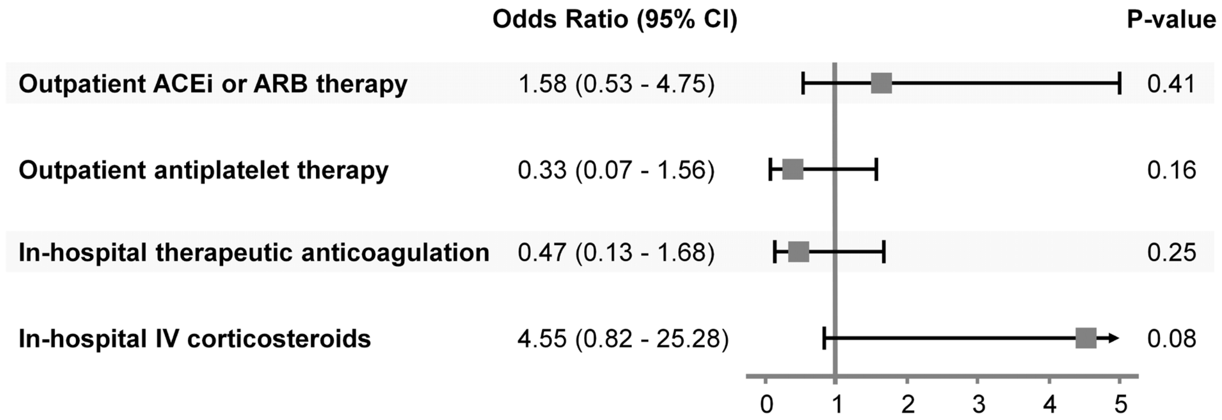

**Figure S2: Principal components analysis of sample level transcript abundance.** Transcript abundance data were collapsed by sample in order to generate a "pseudo-bulk" RNA sequencing dataset. The first two principal components derived from comparison of these data are shown below, where PC1 discriminates the presence of microthrombi, while PC2 separates samples based upon COVID-19 infection.

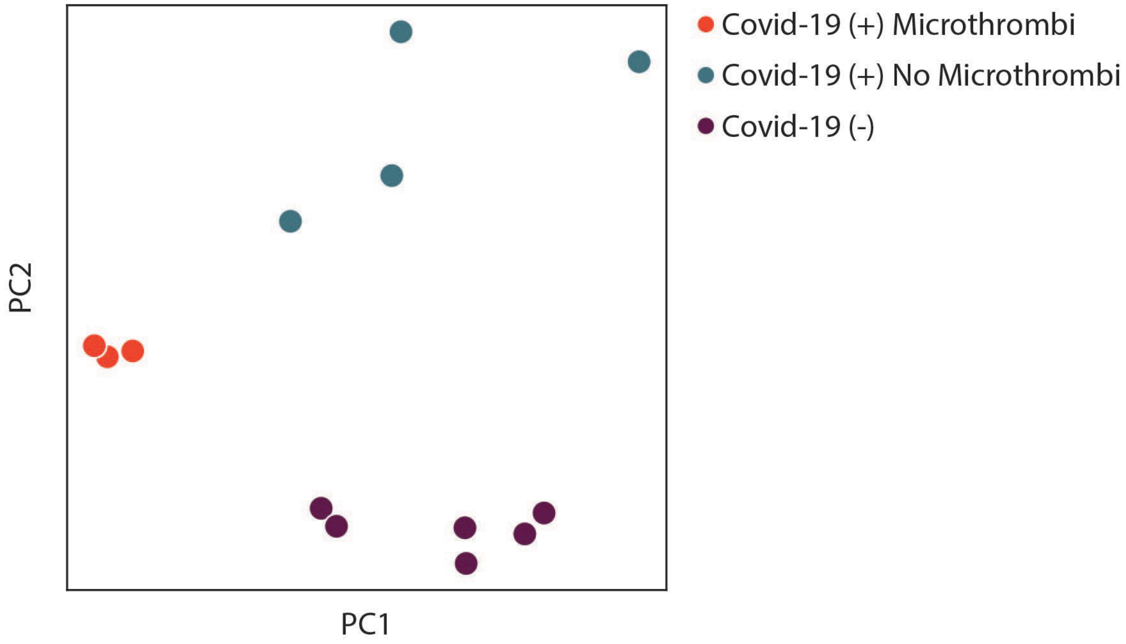

**Figure S3: Identification and removal of low-quality cells in snRNAseq data.** Low quality cells were identified according to mitochondrial gene count, high ratio of exonic to intronic mapping reads and doublet score. Cells flagged in yellow were removed from the data matrix with the resulting post QC UMAP displayed on the right. These filtered data were used for all downstream analyses.

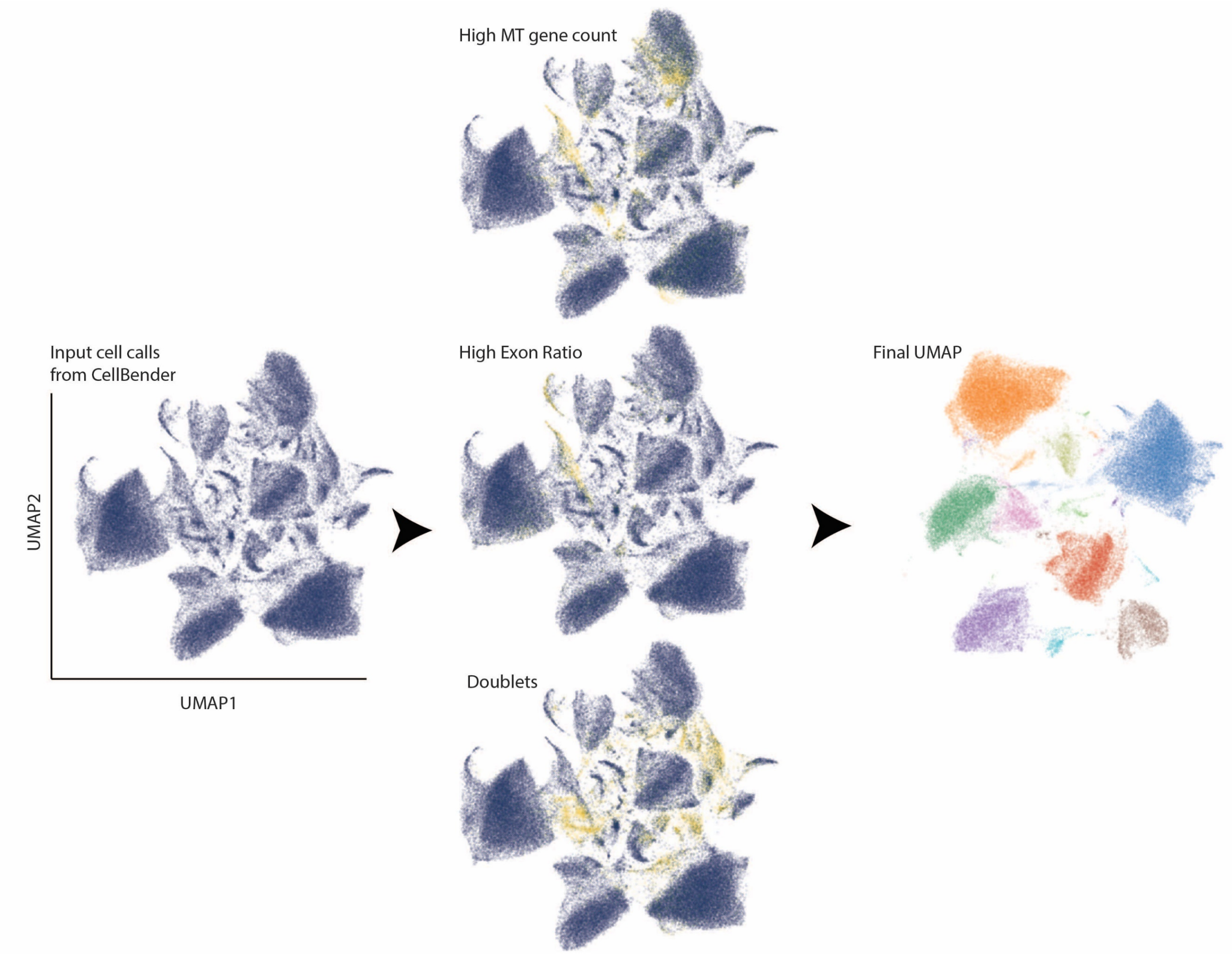

**Figure S4: Volcano plots displaying differential expression results for presence of COVID-19 and for presence of microthrombi.**

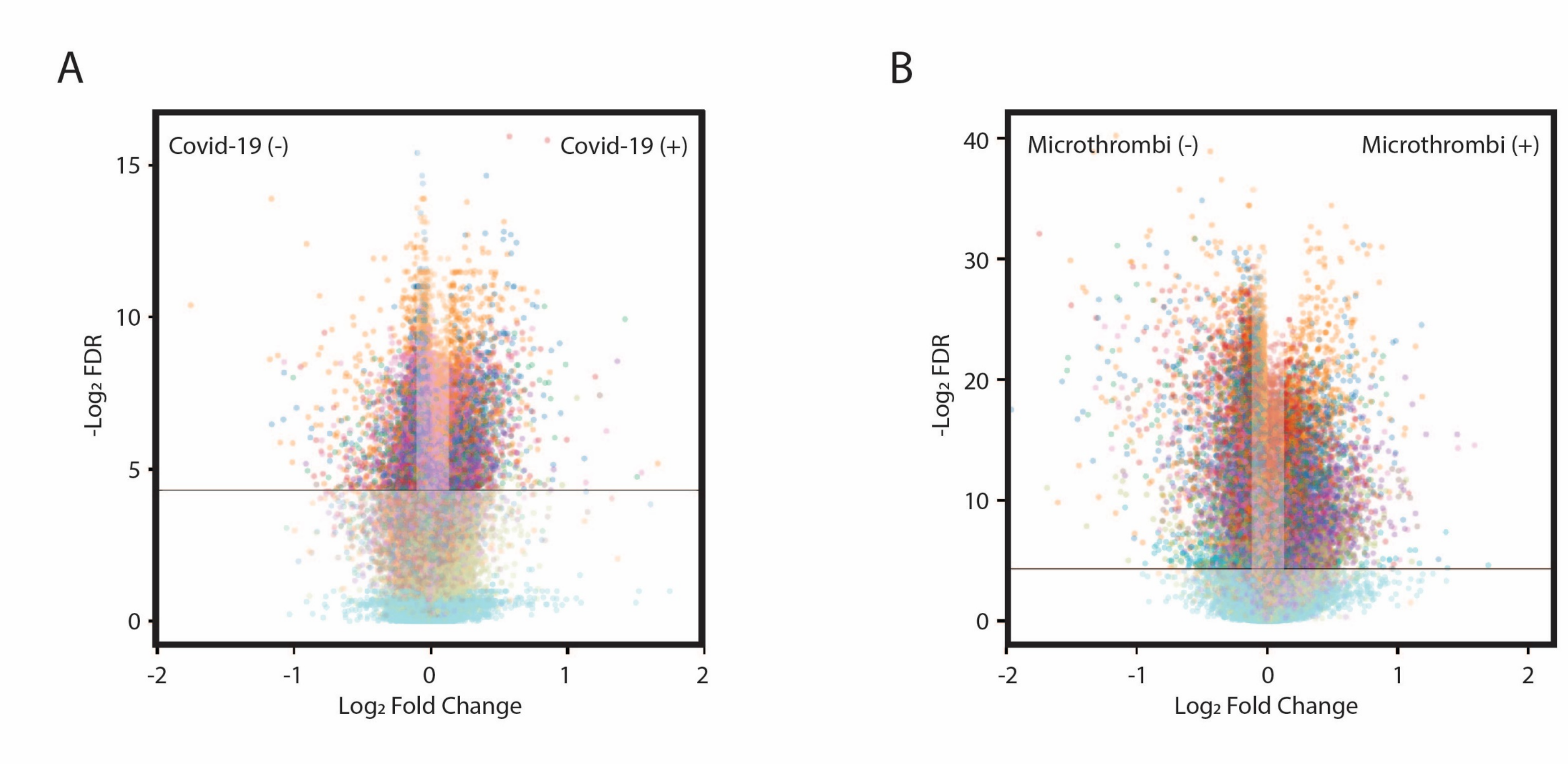

**Figure S5: Genewalk regulators for COVID-19 (+) versus COVID-19 (-) reference samples.**

Regulator genes for each cell type which drive ontology differences when comparing COVID-19 (+) to COVID-19 (-) reference samples. Color corresponds to the cell types as displayed in Figure 3 of the main text.

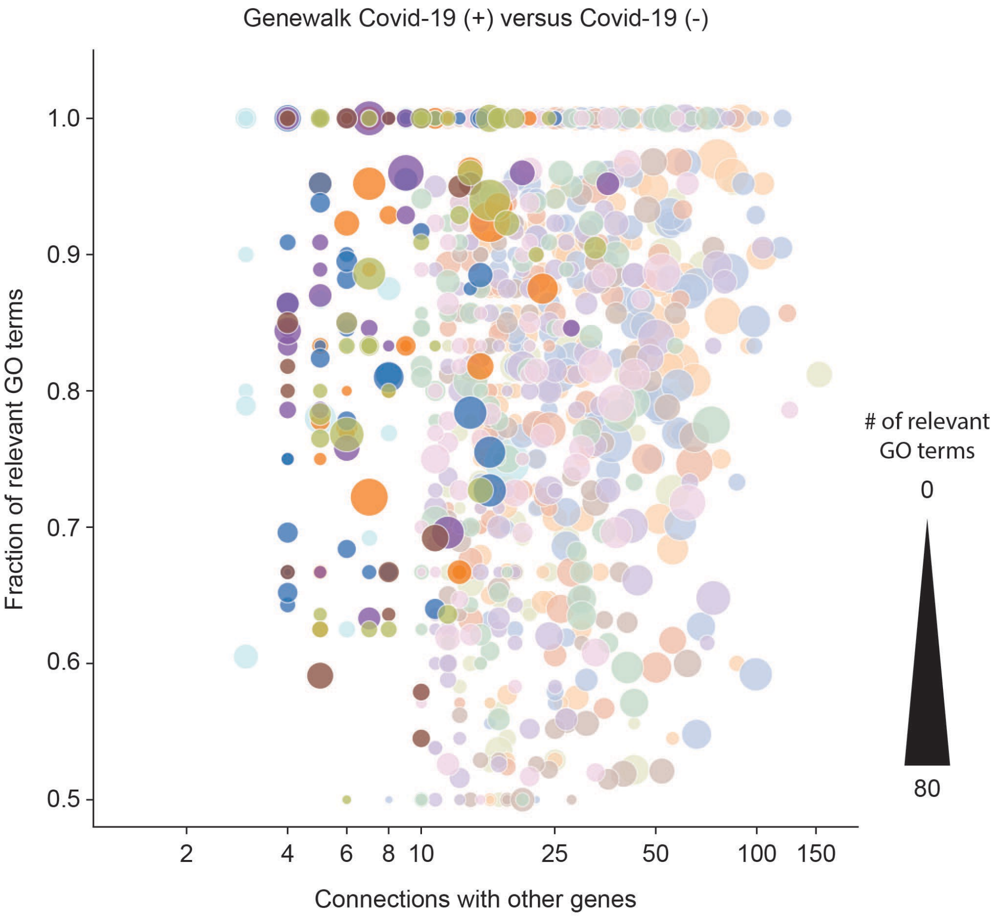

**Figure S6: Ontology analysis for select fibroblast regulator genes identified by Genewalk.**

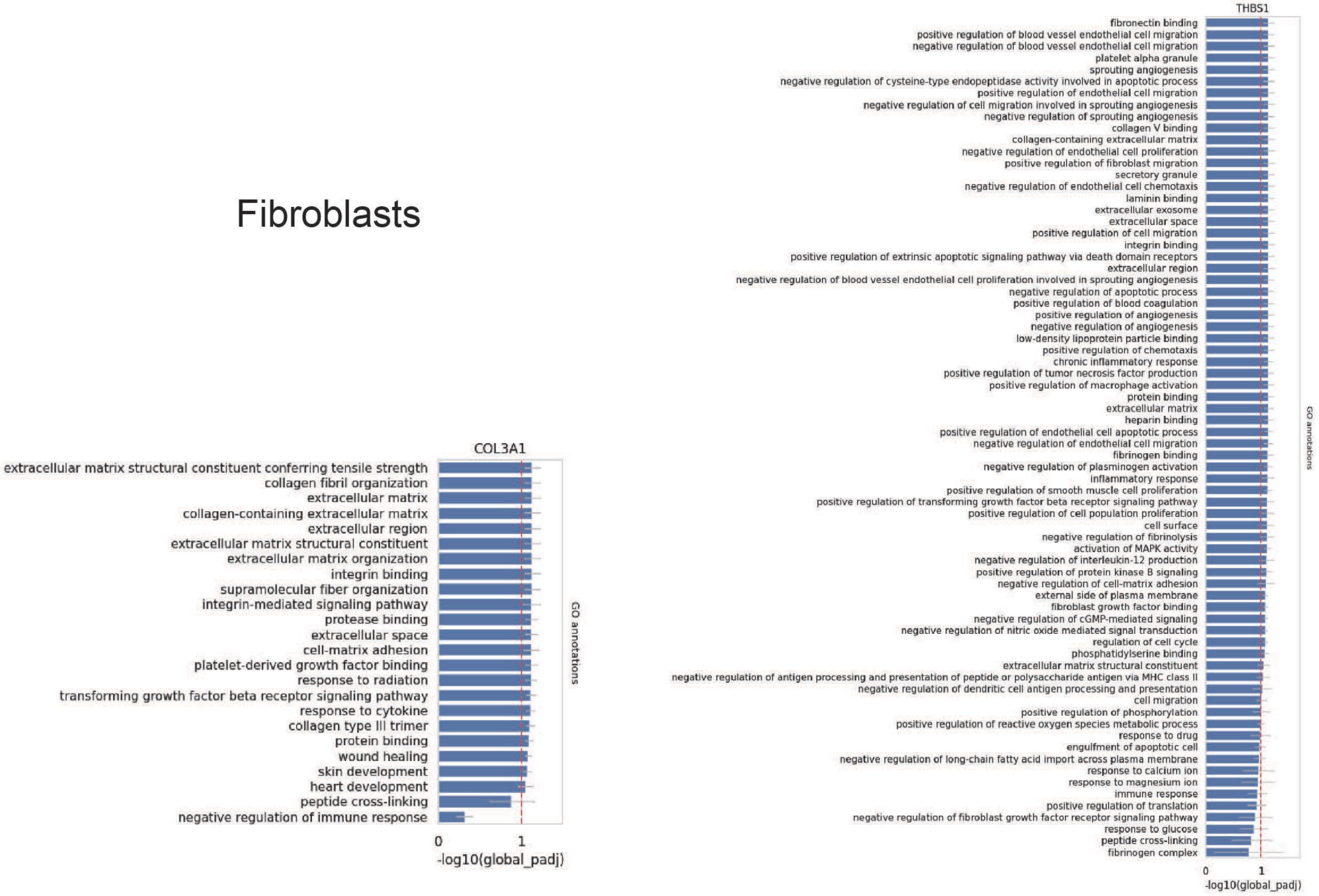
